# Temporal structure in sensorimotor variability: a stable trait, but what for?

**DOI:** 10.1101/817916

**Authors:** Marlou Nadine Perquin, Marieke K. van Vugt, Craig Hedge, Aline Bompas

**Author notes:** Corresponding author: Marlou Perquin, Arbeitseinheit 14 Biopsychologie & Kognitive Neurowissenschaften Universität Bielefeld, Universitätsstraße 25, 33615 Bielefeld Germany.

## Abstract

Human performance shows substantial endogenous variability over time, and this variability is a robust marker of individual differences. Of growing interest to psychologists is the realisation that variability is not fully random, but often exhibits temporal dependencies. However, their measurement and interpretation come with several controversies. Furthermore, their potential benefit for studying individual differences in healthy and clinical populations remains unclear. Here we gather new and archival datasets featuring 11 sensorimotor and cognitive tasks across 526 participants, to examine individual differences in temporal structures. We first investigate intra-individual repeatability of the most common measures of temporal structures – to test their potential for capturing stable individual differences. Secondly, we examine inter-individual differences in these measures using: 1) task performance assessed from the same data, 2) meta-cognitive ratings of on-taskness from thought probes occasionally presented throughout the task, and 3) self-assessed attention-deficit related traits. Across all datasets, autocorrelation at lag 1 and Power Spectra Density slope showed high intra-individual repeatability across sessions and correlated with task performance. The Detrended Fluctuation Analysis slope showed the same pattern, but less reliably. The long-term component (d) of the ARFIMA(1,d,1) model showed poor repeatability and no correlation to performance. Overall, these measures failed to show external validity when correlated with either mean subjective attentional state or self-assessed traits between participants. Thus, some measures of serial dependencies may be stable individual traits, but their usefulness in capturing individual differences in other constructs typically associated with variability in performance seems limited. We conclude with comprehensive recommendations for researchers.

## Introduction

For any action that one repeatedly executes over time, different iterations will show a large amount of variability in their time of execution. Such ‘intra-individual variability’ – variability within the same individual over time – manifests itself prominently during cognitive testing, as participants are commonly instructed to repeat the same actions over a large number of trials. Even in very simple reaction time (RT) tasks, participants’ performance over the trials shows large fluctuations over time (see Figure 1A, top-left panel for an example of the RT series from one participant over 1000 trials). It is clear though that variability reflects more than measurement noise: it is a stable individual trait that transfers across tasks and modalities (Hultsch, MacDonald & Dixon, 2002; Hultsch, MacDonald, Hunter, Levy-Bencheton & Strauss, 2000; Saville et al., 2011; 2012), evolves with age and neurodegenerative disorders (e.g., Tales et al., 2012; Tse, Balota, Yap, Duchek & MacCabe, 2010), and is considered a robust marker of Attention-Deficit and/or Hyperactivity Disorder (ADHD; see Kofler et al., 2013 for a meta-analysis; see Tamm et al., 2012 for a review). Variability has thus proven itself to be a strong candidate for studying individual differences across various neuro-cognitive disciplines.

**Figure 1.**
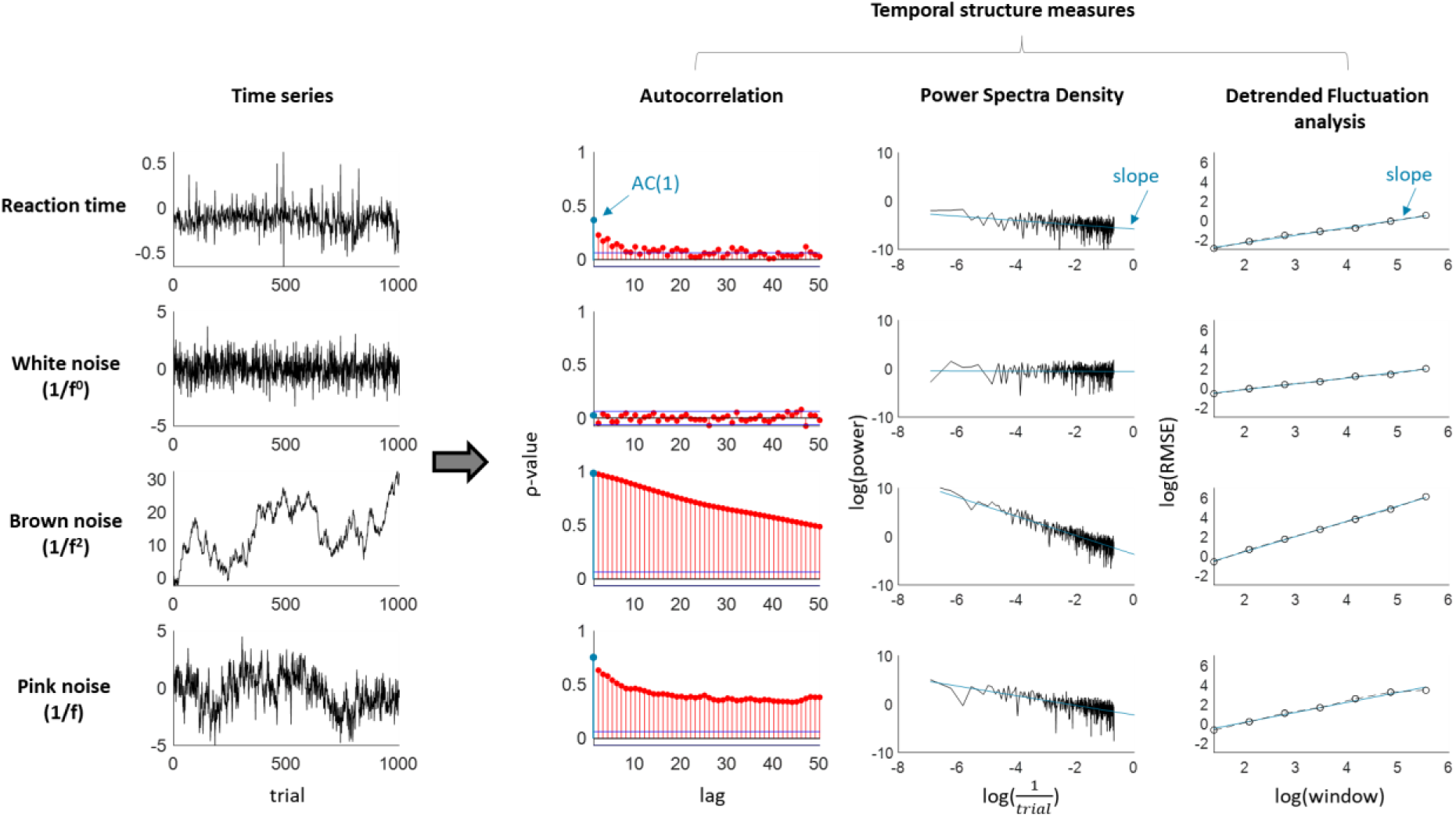
Examples of time series data over 1000 samples (left) and their corresponding temporal structures from lag 0 to lag 50. Shown are a reaction time series (showing small but clear temporal dependency), a white noise series (no temporal dependency), a brown noise series (i.e., a random walk, very high temporal dependency), and pink noise (high temporal dependency with slow decay). The extracted temporal dependency measures (shown in blue) are the autocorrelation at lag 1, the slope of the PSD, and the slope of the DFA.

In experimental data, RT variability is often measured by calculating the standard deviation (SD) or coefficient of variability (SD divided by mean) over whole sessions (e.g., Perquin et al., 2020), but these measures only reflect the overall amount of variance in the data. Instead, quantifying the correlation of an RT series with itself across trials (i.e., autocorrelation) reveals that RT is indeed not independent over time, but often shows a dependency with itself that persist over many trials (see for instance Gilden, 2001; Torre, Vergotte, Viel, Perrey & Dupeyron, 2019; Torre & Wagenmakers, 2009; Van Orden, Holden & Turvey, 2003; Wagenmakers, Farrell & Ratcliff, 2004). This means that the RT on trial *n* is more similar to trial *n+1* or to trial *n+8* than to trial *n+300*, for example. Such dependency is also revealed in plots featuring RT series from a single participant across trials, which do not show a completely random pattern of data points, but rather show (noisy) clusters of fast and slow performance. It may seem probable that quantifications of the trial-to-trial dynamics can give additional information beyond ‘simple’ variability measures.

Hinting towards the fruitfulness of this approach, it has been suggested that these temporal structures differ between individuals (Gilden & Hancock, 2007; Madison, 2004; Simola, Zhigalov, Morales-Muñoz, Palva & Palva, 2017; Torre, Balasubramaniam, Rheaume, Lemoine & Zalaznik, 2011), but this has not been investigated systematically. More specifically, it has been proposed that they reflect the ability for the brain to flexibly adapt and should be a marker of brain health (Simola et al., 2017; Torre et al., 2019). Related to this (although not in direct support of this claim), the temporal structures of *brain activity* (e.g., oscillatory power at rest) have been shown to be altered compared to healthy controls across different neuropsychiatric diseases, including schizophrenia, epilepsy, depression, Alzheimer’s disease, Parkinson’s disease, and autism (see Zimmern, 2020, for an overview). For example, the temporal structures of oscillatory power at rest have been shown to be reduced in schizophrenia (Golnoush et al., 2020; Nikulin, Jönsson & Brismar, 2012; Slezin et al., 2009; Sun et al., 2014; but see Cruz et al., 2020) and epilepsy patients (Adda & Benoudnine, 2016), and increased in depression patients (Gärtner et al., 2017), compared to healthy controls.

However, temporal structures also come with controversies regarding the best way to measure them, their time scale, their origins and interpretation. It has been argued they might reflect switches in mental states or mental flexibility, but empirical evidence for this claim is lacking. It is also unknown if the temporal structures show intra-individual reliability over sessions and tasks – a necessary requirement for them to be useful biomarkers and help us characterising individual differences (Hedge, Bompas & Sumner, 2020; Hedge, Powell & Sumner, 2018; Mayeux, 2004). In the current study, we conduct a large-scale analysis of temporal structures in RT series across various sensorimotor and cognitive tasks from new and archival datasets. First, we quantify short- and long-term structures in RT series across different tasks. Next, we test the repeatability of temporal structures over sessions, and across tasks – to examine the extent to which the structures can be used to study (stable) individual differences. In the third part, we test if individual differences in temporal structures show any relationship to other constructs related to behavioural variability, elaborated on in the following section.

### Fluctuations in behaviour and attentional state

The causes of variability in performance remain largely mysterious, attracting both physiological and cognitive interpretations. Intuitively, we are quick to associate fluctuations in our behaviour with fluctuations in our attentional state – e.g., “I was too slow because I was not focused on my task”. Indeed, variability is often thought to be causally related to occasional disengagement from the task (Anderson, Petranker, Lin & Farb; 2021; Laflamme, Seli & Smilek, 2018; Seli, Cheyne & Smilek, 2013; Thomson, Seli, Besner & Smilek, 2014), and both are known to be increased in ADHD patients (ADHD and off-taskness: Shaw & Giambra, 1993; ADHD and variability: Kofler et al., 2013; Tamm et al., 2012). As our attentional state waxes and wanes over time between being on- and off-task, it makes sense that the fluctuations in performance are not random, but likewise temporally structured.

Given the intuitive link between temporal structure and attentional state, one may specifically expect individual differences in the strength of serial dependencies to be informative of people’s ability to sustain attention and high performance over time. In the present research, we therefore link individual differences in variability, and more specifically temporal dependencies, with subjective ratings of on-taskness and self-reported traits associated with attention deficits.

Before tackling this, let us first introduce to less familiar readers what temporal dependencies are about, and review the main methods used to quantify them. To aid interpretation, we compare the effect of each measure on white, brown, and pink noise – each of which is characterised by its own typical time structure, as well as on example RT series (see Figure 1 for an overview). Readers familiar with these concepts may go straight to the next sections “*Why might temporal structures be interesting?*” and *“Individual differences in temporal dependencies*”. We then introduce our novel study, as well as archival datasets brought-in to strengthen and broaden specific aspects of our conclusions.

### Temporal structures and quantification

One difficulty in navigating the literature is the variety of measures used by researchers, some simple, others more complex. These are introduced briefly here (details on all four quantification methods can be found in the Methods section).

Autocorrelations are the most straightforward characterisation of temporal dependency and can be estimated for any delay (Box, Jenkins, Reinsel & Ljung, 2016; with delay referring to trials in experimental RT series). RT data typically show positive autocorrelations at short lags (Figure 1, top row). This is inconsistent with a completely random process in which the observations are fully independent from each other (i.e., white noise), and which would show no autocorrelation at any lag (Figure 1, second row). In contrast, a process in which each observation is just the combination of the preceding observation *n-1* plus random error (i.e., ‘brown noise’), the autocorrelation at lag 1 (AC1) is high (near one) and shows a very slow decay over the subsequent lags – theoretically never reaching zero.

Most studies use more complex methods to study temporal dependencies, the most common being the Power Spectrum Density (PSD; Box et al., 2016) and Detrended Fluctuation Analysis (DFA; Peng, Havlin, Stanley, & Goldberger, 1995). For the PSD, the RT series is analysed in the frequency domain through Fourier-transform (Box et al., 2016), and a regression line is fitted between the inverse of trial number and the power spectrum, after both have been log-transformed. The slope of this regression line represents the amount of temporal structure: a slope of 0 indicates the absence of any structure (white noise with an SD of 1), while brown noise series result in a slope of 2 (Figure 1). For DFA, the entire series is divided into windows, which size and amount of overlap is determined by the researcher. The datapoints in these windows are detrended and reduced to an ‘average fluctuation’, that reflects the distance of each data point to the trend line. Similar to the PSD, the linear slope fitted in the log-log space of window size and average fluctuation (Peng et al., 1995) represents the temporal structure. A white noise series with an SD of 1 will be reflected in slope *α* = .5, brown noise in *α* = ~2-3, and anticorrelated series in *α* < .5. Rather than giving an estimate for each delay separately, these methods thus provide one parameter reflecting the degree of temporal dependency across the entire RT series.

While white noise shows no temporal dependency and brown noise shows high temporal dependencies, pink noise lies in-between the two. Pink noise is also known as ‘1/f noise’, with its power being equal to 1 over frequency (Figure 1; although anything between ‘1/f^.5^ to ‘1/f^1.5^ noise’ still typically considered as 1/f noise). It is characterised by relatively high autocorrelation at short lags, which slowly but gradually decreases to zero over the larger lags. Pink noise is an important concept within the literature on temporal structure, as it has been claimed to best match behavioural time series. We come back to this in the “Criticality” section below.

Although PSD and DFA have been popular for analysing RT, the methods have an important limitation: they can be ambiguous about what drives the observation of temporal dependency: merely the correlation between trials close together (‘short-term dependency’), or (also) the correlation between more distant trials (‘long-term dependency’). As pointed out in prior literature (see Wagenmakers et al., 2004 for a first detailed exposition of the problem), non-zero (or for DFA, non-.5) slopes are not necessarily indicative of long-term dependencies. Indeed, although short-term dependencies should theoretically lead to shallower slopes, in practice they can resemble pink noise. To solve this ambiguity, the use of autoregressive fractionally integrated moving-average (ARFIMA) models has been suggested, which can explicitly test the necessity of a long-term dependency parameter over short-term parameters only (Wagenmakers et al., 2004; Torre et al., 2007).

### Why might temporal structures be interesting?

#### Attentional state

Just as our behaviour shows fluctuations over time, so do our meta-cognitive states. For example, throughout a task, we may feel more on-task on some moments and more off-task on others. It has been found that these fluctuations in subjective attentional state correlate locally with fluctuations in performance (e.g., consistency during synchronised tapping), which deteriorates when one feels more off-task (Anderson et al., 2021; Laflamme et al., 2018; Seli et al., 2013; Thomson et al., 2014). These findings seem to match common intuitions about our own functioning – namely, that we may show streaks of good performance during which we feel extremely focused as well as streaks of poor performance in which we feel less on-task (e.g., Gilden & Wilson, 1995; Smith, 2003). It has been argued that increased fluctuations from on-taskness to off-taskness are reflected in increased temporal structures (Irrmischer, van der Wal, Mansvelder, & Linkenkaer-Hansen, 2018). Indeed, a mechanistic model has been proposed, where the combination of short-term dependencies (first order autoregressive term) and (comparatively slower) alternation between two response modes or strategies suffice to capture empirically observed temporal dependencies in synchronised tapping (Torre & Delignières, 2008; Torre & Wagenmakers, 2009; Torre, Balasubramaniam & Delignières, 2010, see also Bastian & Sackur, 2013). Although it is tempting to associate the two modes to subjective judgments of being on-task and off-task, note that they can also be conceived as two states of an internal parameter (e.g., response threshold) which may not have meta-cognitive counterparts.

If temporal structures are indeed related to the pace at which such an internal variable fluctuates, they may be different in people who show low consistency in task performance. ADHD has previously been associated with higher RT variability, which has been attributed to more attentional lapses, but also with a lack of response inhibition, the combination of which may lead to a pattern of extremely slow and extremely fast responses (Kofler et al., 2013; Tamm et al., 2012). Some previous work has examined temporal structures in performance of ADHD patients (e.g., Castellanos et al., 2005; Geurts et al., 2008; Johnson et al., 2007; see Karalunas, Huang-Pollock & Nigg, 2012; Karalunas, Geurts, Konrad, Bender & Nigg, 2014 for reviews; see Kofler et al., 2013 for a meta-analysis). While these studies do not report the PSD slope (nor any of the time series analyses mentioned above), they found increased power in the low frequency of the spectrum (< 1.5 Hz) in ADHD patients (although it is unclear whether these would translate into higher PSD slopes for ADHD patients; see Discussion more details).

#### Criticality

One common reason why the existence of temporal structures in behaviour has piqued interest is because they may provide fundamental insights into how cognition emerges from dynamical brain systems. Most commonly, they have been studied in the framework of criticality – though of course, they could be informative of our functioning regardless of its link with one particular framework.

In short, critical systems are thought to reflect an optimal balance between predictability and randomness. In physics, a system that has converged to the border between order and chaos is called critical, and natural fluctuations within such metastable systems exhibit a 1/ƒ spectrum (Thornton & Gilden, 2005). It has been argued that neural networks self-organise to operate around the critical point, giving them maximum sensitivity to perturbation (e.g., from sensory inputs) without activity imploding (see e.g., Beggs & Timme, 2012; Shew & Plenz, 2013 for detailed reviews). To the extent that these theories apply to human behaviour and cognition (note the big conceptual jump), the presence of 1/f noise in behavioural time series could be taken to imply that 1) the brain-body system approaches criticality and 2) the system is affected by very slow fluctuations that “cause a cascade of energy dissipation at all length scales” (Bak, Tang & Wiesenfeld, 1987). This cascading means that critical systems display some amount of correlation (here we focus on temporal correlations), with trials close in time showing the strongest correlation, which decreases as the time lag increases. As the dependency over time is neither perfect nor random, this is most similar to the pink noise described above. Indeed, critical systems are thought to show such pink noise. Within the literature, this is often referred to as a ‘power law’ – the PSD shows up as a constant negative slope throughout. This line reflects that the relationship is ‘scale-free’: One can take any subpart of the spectrum and find the same straight line; it has no specific time scales. Although not all critical systems adhere to the power law, and power laws can show up in non-critical systems, it is generally seen as a highly important characteristic of critical systems.

Behaviour over time also shows 1/f noise and it has therefore been argued that cognition is a self-organised critical system (e.g., Gilden, 2001; Kello, Beltz, Holden & Van Orden, 2007; Thorston & Gilden, 2005; Van Orden et al., 2003). However, whether or not the magnitudes of time structures are actually high enough to be considered pink noise or whether they provide the best fit to the temporal dynamics in behaviour remain controversial topics (see Farrell, Wagenmakers & Ratcliff, 2006; Pressing & Jolley-Rogers, 1997; Wagenmakers et al., 2004; 2005; Wagenmakers, van der Maas & Farrell, 2012 for critiques) – and may be dependent on the analysis method. Nonetheless, the interest in human cognition as a critical system partly explains why the literature has mainly focused on measuring temporal structure (as opposed to manipulating it, or examining its individual differences): The interest often starts and stops at the mere existence of pink noise in the data, focusing on the ‘ubiquitousness’ of this phenomenon, as a main concern of the framework lies with generality across fields (e.g., physics, economics, biology) rather than with finding underlying neuro-cognitive processes (Wagenmakers et al., 2012). As such, temporal structure has, among other examples, been found in simple RT (Wagenmakers et al., 2004; Van Orden et al., 2003), choice RT (Kelly, Heathcote, Heath & Longstaff, 2001; Wagenmakers et al., 2004), mental rotation (Gilden et al., 2001), visual search, lexical decision, word naming, shape discrimination, and colour discrimination (Gilden, 2001; Van Orden et al., 2003), go/no-go (Simola et al., 2017), racial implicit bias tasks (Correll, 2008; Maduski & LeBel, 2015), and speech (Kello et al., 2008) – although evidence for the non-universality of temporal structure has also been found previously (see Wagenmakers et al., 2004 for an overview). Its existence is particularly clear in specific tasks, including the task used in the present article: finger tapping in synchrony with a tone (Torre et al., 2010). The present research therefore addresses the usefulness of this phenomenon in understanding individual differences.

#### Predicting behaviour

Even if the temporal structures do not relate to criticality or attentional state, one may still agree that RT series carry a predictable component—carried by these measures— and an error component. However, while prior studies have used time series analyses to quantify the structure in an existing series, it remains unknown to what extent these structures are informative for future behaviour. In other words, if one can find that behaviour on trial *n* is correlated to trial *n-1*, is it also possible to predict behaviour on yet-unobserved trial *n+1*?

Such ‘forecasting’ lies within the possibilities of the time series analysis, particularly of the ARFIMA models. These have been used to forecast weather or economic trends, and a recent article outlines a method to assess the feasibility of this approach to forecasting human behaviour on the next trial (Wagenmakers, Grünwald & Steyvers, 2006). This may be complementary to calls to make psychology a more predictive science in order to better understand human behaviour (Yarkoni & Westfall, 2017). Aside from a theoretical interest, such behavioural forecasting may also have a practical use: Given that real-life behaviour also fluctuates over time and occasionally fluctuates to extremely poor responses (car accidents would be real-life equivalents of very long RT, errors, or omissions), it would be desirable to prevent these poor responses by predicting them before they occur, based on past behaviour. Of course, the fruitfulness of this approach is dependent on the existence of temporal structures.

### Individual differences in temporal dependencies

A few recent studies have reported weak to moderate correlations between DFA slopes and individual differences in performance during cognitive tasks. First, Smit, Linkenkaer-Hansen & de Geus (2013) reported a negative correlation in a tapping task involving pressing a key every second without an auditory reference – indicating that a high DFA slope (i.e., more temporal dependency) was associated with poorer task performance. Irrmischer et al. (2018) also found a negative correlation with performance in a sustained attention task, as measured by RT to rare target stimuli. In a second study using the same task, RT and slopes were higher after negative mood induction (thought to increase mind wandering, Smallwood, Fitzgerald, Miles & Philips, 2009) compared to positive but not to neutral mood induction (note that the study did not include a pre-manipulation measure of the task). In contrast, Simola et al. (2017) reported a positive correlation with performance in the Go/No-Go task – indicating more temporal dependency was associated with fewer commission errors – with no correlation with mean RT or standard deviation of RT.

Despite the varied findings, their interpretations rely on the same theoretical viewpoint: brains which operate closer to the critical point show higher long-term correlations. While Simola et al. (2017) take their positive correlations as evidence that criticality allows for the mental flexibility demanded by some tasks, Irrmischer et al. (2018) interpret their negative correlations as evidence that criticality allows for the successful dynamics of switching attention from task-related to task-unrelated thoughts on their sustained attention task. We come back to the role of task demands in the Discussion.

Few studies have looked at the intra-individual reliability of temporal dependency in task performance, both within and across sessions and tasks. Smit et al. (2013) observed poor to moderate split-half reliability of the PSD and DFA slopes. Torre et al. (2011) reported moderate repeatability of the DFA slopes on two tasks (a circle drawing and a tapping task), but found no cross-task correlations. This relative stability of temporal structures in behavioural performance over time is consistent with the stability observed in neural oscillations (Nikulun & Brismar, 2004; Smit et al., 2013; see Discussion for more details).

### Current research

Here, we examine individual differences in temporal dependencies, to assess: 1) to what extent these structures repeat in individuals over time, 2) to what extent these structures repeat in individuals across different tasks, 3) how these structures relate to objective and subjective task measures, and 4) how these structures relate to self-assessed attention-deficit related traits. Prior to investigating these questions, we first verify the presence of the temporal structures in our data, as this is a necessary condition for examining any individual differences. We analysed data from two cohorts which we specifically collected for the current project, as well as archival datasets previously collected for other purposes. Our study used the Metronome Response Task (MRT; Seli et al., 2013), in which participants are instructed to press a button in synchrony with a regular tone. Throughout the MRT, participants are pseudo-randomly presented with thought probes asking them to judge their attentional state. This task comes with several benefits for our current interests.

First, the behavioural task relies minimally on the external environment while still providing a behavioural measure and is therefore particularly suited to assess *endogenous* fluctuations in performance. The rhythmic reaction time provided on each trial offers continuous access to fast fluctuations in the underlying cognitive functions (as opposed to tasks featuring accuracy scores, where a continuous performance measure can only be obtained over multiple trials that are each much longer than the responses in the MRT). Secondly, the MRT also provides an *online* measure of attentional state (i.e., measured during the experiment), which is known to correlate locally to fluctuations in RT. Thirdly, tapping- and time-estimation based tasks (both with and without metronome) have been used extensively in the motor literature and show clear temporal structures (e.g., Chen, Repp & Patel, 2002; Delignières, Lemoine & Torre, 2004; Ding, Chen & Kelso, 2002; Gilden, Thornton & Mallon, 1995; Lemoine, Torre & Delignières, 2009; Madison, 2001; Wagenmakers et al., 2004). Last, the main performance measure is straightforward to interpret, in contrast to tasks such as Go/No-Go, that require both withholding and responding, and provide multiple measures of performance, such as ‘omission errors’, ‘commission errors’, and ‘RT to target stimuli’. To get a full picture of performance, these different performance measures must be interpreted together to take into account factors such as speed-accuracy trade-offs. For instance, if a participant produces only few commission errors (she does not respond when she shouldn’t) but also many omission errors (she also does not respond when she should), it is unclear whether this constitutes good or poor performance. This complexity allows arguably too much flexibility in results interpretation. In contrast, the metronome task produces one main measure (asynchronies, or Rhythmic RT) with only very few omissions (<1% for most participants), that can be safely ignored. All in all, this means that, if one does not find consistency and external validity of temporal structures in MRT performance, it is unlikely one would find it on other tasks.

To generalise our result patterns, we also analysed three archival datasets. The first dataset contains behavioural data from seven tasks, which were collected for a study on intra-individual reliability of task performance in different tasks related to cognitive control (Hedge, Powell & Sumner, 2018). This includes the data of 104 participants who performed two sessions of the Eriksen Flanker, Stroop, Go/No-go, and Stop-signal tasks, and 40 participants who performed two sessions of the NAVON, SNARC, and Posner tasks. The second dataset contains behavioural data from a SART and a Visual Search task (30 participants). This dataset was collected for a study on EEG markers of subjective attentional states, both within and across tasks (Jin, Borst & van Vugt, 2019) – and thus contains subjective ratings from occasionally presented thought probes. The third dataset also uses the MRT (Anderson et al., 2021) and consists of a large-N study (N = 375) investigating the reproducibility of previously reported relationships between subjective attentional states and behavioural variability, intentionality, and motivation. None of these are a perfect match for our aims, but each provides valuable contributions in a different way.

We present our results in three parts (see Table 1 for an overview). In the first part of the results section, we validate the existence of temporal dependencies, including long-range correlations, in the individual data series of each of these tasks. The second part relates to the within-subject repeatability of temporal structures (across different time points and different paradigms), and the third part relates to their potential between-subject correlations with performance and metacognitive attentional state ratings. Below we elaborate on these research aims, and specify which datasets are suited for which particular questions.

**Table 1.** Overview of our current research questions and corresponding methods, and the datasets used to answer each question.

| Research question | Method | Datasets |
| --- | --- | --- |
| *Part I. Validation* | - One-sample t-tests on distributions of | New data |
| Do RT series show short- and long-term temporal structures? | temporal structure measures to test against | Anderson et al., 2021 |
|  | null-hypothesis of no structure | Jin et al., 2019 |
|  | - ARMA/ARFIMA modelling to test for | Hedge et al., 2018 |
|  | presence of long-term structure | Wagenmakers et al., 2004 |
| *Part II. Intra-individual repeatability* | Pairwise correlation analyses |  |
| A. Are temporal structures repeatable over time? |  | New data |
| B. Are temporal structures generalisable across tasks? |  | Jin et al., 2019 |
|  |  | Hedge et al., 2018 |
| *Part III. Inter-individual differences* | Pairwise correlation analyses |  |
| A. Do temporal structures correlate with task performance |  | New data |
| calculated on the same data series? |  | Anderson et al., 2021 |
| B. Do temporal structures correlate with subjective ratings on |  | New data |
| attentional state measured throughout the same task? |  | Anderson et al., 2021 |
| C. Do temporal structures correlate with self-assessed traits from |  | New data |
| attention-deficit related questionnaires? |  | Anderson et al., 2021 |

#### Part I. Validating the presence of temporal dependencies

Separately for each dataset (and where relevant, each session), the temporal dependency measures (AC1, PSD slope, DFA slope, and ARFIMA parameters) were quantified on each participant’s RT series, and statistically compared to chance. Furthermore, fit values from the ARMA and ARFIMA models were compared for each participant, to specifically validate the presence of *long-term* dependency.

#### Part II. Intra-individual repeatability of temporal dependency

For temporal dependencies to be informative for any individual differences, they need to be a stable trait, i.e., show consistency within individuals. We report the intra-individual repeatability of the temporal dependency measures (AC1, PSD slope, DFA slope, and ARFIMA parameters) over two sessions of the MRT (conducted about 30-50 minutes apart). The measures were calculated separately for each session (time 1 vs time 2). For comparison, the same analysis was applied to task performance (as measured by behavioural variability) and subjective attentional state ratings.

To anticipate our results, we indeed found the temporal structures in MRT data to be repeatable over time. These findings beg the question to what extent stable temporal dependencies are found in tasks that are used more commonly in the neuro-cognitive literature. Therefore, we also examined the intra-individual repeatability of the temporal dependency measures in RT series of a diverse set of well-established cognitive control/impulsivity related tasks (Hedge et al., 2018).

As a next step, we examined if the temporal dependencies are also stable across different tasks – i.e., if the temporal dependency of an individual on one task is informative for their temporal dependency on another task. For this question, we used the seven cognitive tasks as well as a SART and Visual Search data (Jin et al., 2019) of a different dataset — both cases in which the same participants performed multiple tasks. Temporal dependency was correlated between subjects across the cognitive tasks (i.e., across each pair of the Eriksen Flanker, Stroop, Go/No-go, and Stop-signal tasks, and across each pair of the NAVON, SNARC, and Posner tasks), as well as across RT series in a SART and Visual Search task (30 participants).

#### Part III. Between-subject correlates of temporal dependency

In the third part, we report the extent to which temporal structures derived from RT series relate to individual differences in: 1) task performance measures calculated on the same data, 2) subjective reports of attentional state from thought probes, and 3) self-assessed personality traits from attention-deficit related questionnaires. For these between-subject analyses, we always use measures from the first session, for consistency across datasets.

##### Performance

While performance on the MRT task is straightforwardly quantified, performance on the neurocognitive tasks can be computed in many different ways (mean RT or accuracy across all trials, or in each subcondition – congruent or incongruent, or the difference between them etc). To keep the analysis feasible, we therefore used only the MRT data (both our own and Anderson et al.) to examine the relationship between temporal dependency and performance.

##### Subjective attentional state

To examine the between-subject relationship between temporal structure and subjective attentional state, we used all the datasets that contained subjective ratings of such attentional states. These are both MRT datasets, the SART and the Visual Search data.

##### Self-assessed traits

For the relationship between temporal structure and self-assessed attention-deficit related traits, we used both MRT datasets, as these included relevant questionnaires. In our current design, all participants completed a questionnaire on ADHD tendency. As ADHD is a multi-faceted condition, potential correlations between temporal structure and ADHD tendency would reveal little about the driving mechanism. For the first cohort, we therefore also included a questionnaire on impulsivity (one of the main two facets of ADHD) and a questionnaire on mind wandering tendencies, which has been associated with ADHD tendencies both in healthy and clinical participants (Perquin & Bompas, 2019; Shaw & Giambra, 1993; Seli, Smallwood, Cheyne & Smilek, 2015; Unsworth, Robison & Miller, 2019) and may reflect an individual tendency towards getting off-task. However, anticipating on our results, we found clear statistical evidence against between-subject correlations across all three questionnaires and temporal dependency on the first cohort. For the second cohort, we therefore only included the ADHD questionnaire.

Participants from the Anderson et al. (2021) study completed the Attention-Related Cognitive Errors Scale (Cheyne, Carriere, & Smilek, 2006), which aims to measure an individual tendency to make cognitive errors in daily life that are caused by lapses of attention. This questionnaire has been found to positively correlate with ADHD tendencies in healthy participants (Malkovsky, Merrifield, Goldberg & Danckert, 2012). Anderson et al. (2021) found a modest between-subject correlation between the ARCES scores and behavioural variability on the MRT.

## Methods

Here, we report a short summary for the new and archival datasets we have analysed for the current study (see Table 2 for an overview with the key features of each dataset). An extensive description of the methods can be found in the Appendix.

**Table 2.** Overview of the datasets analysed in the current research, showing both our new collected data and the archival datasets. Listed are the source (‘dataset’), the total number of subjects before exclusion (‘# subjects’), the tasks (‘tasks’), the total number of trials for each task (‘# trials’), whether the participants performed the tasks once or twice (‘# sessions’), whether thought probes on attentional state were included, and if so, on what scale (‘thought probes?’), and the questionnaires we analysed in the current study (‘questionnaires’).

| Dataset | # subjects | Tasks | # trials | # sessions | Thought probes? | Questionnaires |
| --- | --- | --- | --- | --- | --- | --- |
| New data | 139 | Metronome Response Task (MRT) | 1050 | 2* | Yes, on scale from 1-9 | Adult ADHD Self-Report Scale (ASRS)<br>Daydreaming Frequency Scale (DFS)<br>UPPS-P Impulsive Behaviour Scale (UPPS-P) |
| Jin et al., 2019 | 30 | Sustained Attention to Response Task (SART) | 378-486 | 1 | Yes, categorical responses |  |
|  |  | Visual Search | 420 |  |  |  |
| Anderson et al., 2021 | 375 | Metronome Response Task | 900 | 1 | Yes, on scale from 1-6 | Attention-Related Cognitive Errors Scale (ARCES) |
| Hedge et al., 2018, Exp. 1 + 2 | 107 | Eriksen Flanker | 720 | 2 | No |  |
|  |  | Stroop | 720 |  |  |  |
|  |  | Stop-signal | 720 |  |  |  |
|  |  | Go/No-Go | 720 |  |  |  |
| Hedge et al, 2018, Exp 3 | 40 | NAVON | 640 | 2 | No |  |
|  |  | Spatial-Numerical Association of Response Codes (SNAR) | 640 |  |  |  |
|  |  | Posner Cuing | 640 |  |  |  |
| Wagenmakers et al., 2004 | 6 | Simple RT (short and long ITI) | 1024 | 1 | No |  |
|  |  | Choice RT (short and long ITI) | 1024 |  |  |  |
|  |  | Fixed temporal estimation (short and long ITI) | 1024 |  |  |  |

\* 73 out of 139 participants were tested for the second session.

### Collected data

The Metronome Task (Seli et al., 2013) was used to obtain a RT series for each participant (Figure 2). From these series, we calculated for each participant: 1) the standard deviation of the RT, reflecting an overall measure of performance on the task, and 2) temporal dependency in the RT series. The MRT also measured participants’ subjective ratings of attentional state quasi-randomly throughout the experiment. Although the original MRT task offered only three levels of responses (“on task”, “tuned out” and “zoned out”), we offered instead a scale from 1 (completely on task) to 9 (completely off task) to get a more gradual response. For participants who performed the MRT twice, these measures were extracted separately for both occasions.

**Figure 2.**
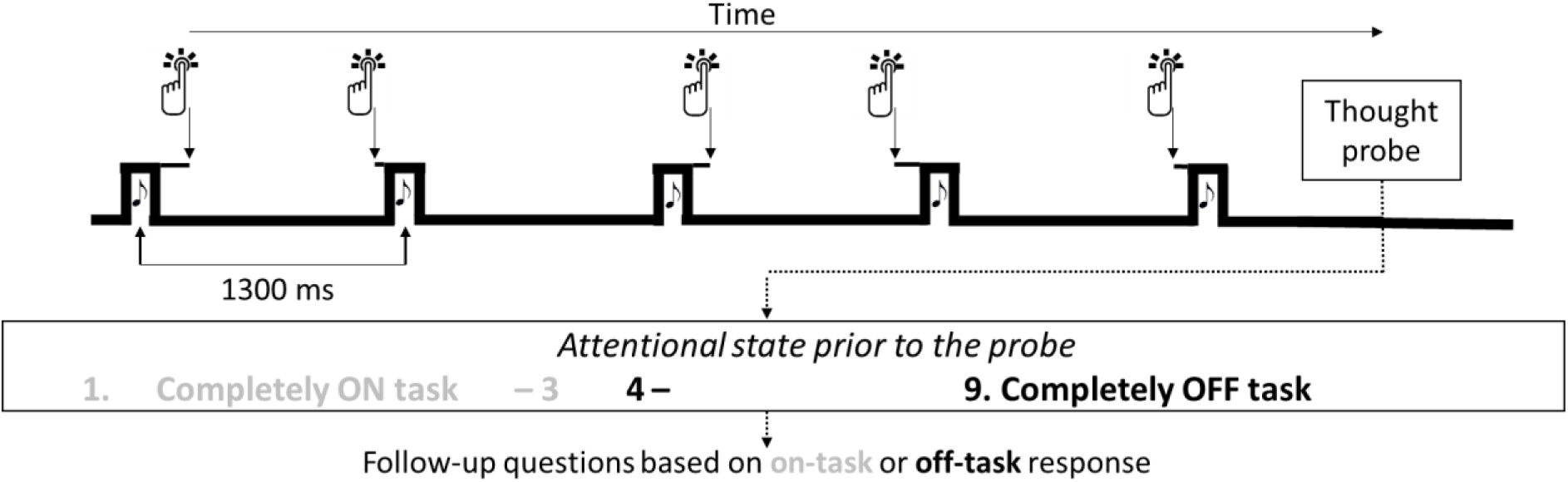
Overview of the task and thought probes.

### Time series analyses

We quantified temporal structures in the RT series using AC1, PSD, DFA, and ARFIMA(1,d,1). Though one might expect that missing data are problematic for time series analyses, the most common method in the literature is to ignore missed responses altogether. In line with this, we excluded omissions from the RT series for all analyses, but verify our results with two alternative imputation approaches (see section *Control analyses*; also see section *Missing data* in the Discussion).

#### Autocorrelation

Autocorrelation quantifies the correlation of a time series with itself over a specified lag (Box et al., 2016). Here, and throughout our analyses, time refers to trials, as typical in the field when dealing with reaction time data series. The autocorrelation *ρ* at lag *k* is given by:

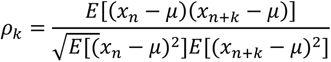

with *x_n_* reflecting observation *n* in time series *x, μ* reflecting the mean of the complete time series, and *E* reflecting the expected value.

We focus on the autocorrelation at lag 1 (correlation of trial *n* with trial *n+1*, AC1), representing the temporal structure at the shortest delay possible in RT series. The AC1 (i.e.,) was calculated for each participant in R (R Core Team, 2013) on their RT series with the *acf* function in the *Forecast* package (Hyndman et al., 2018; Hyndman & Khandakar, 2008).

#### Power Spectral Density

By Fourier-transforming the RT series and calculating the squared amplitude, one can obtain the power spectrum of the series – or equivalently, the power spectrum can be calculated with a Fourier transform on the autocorrelation function (Box et al., 2016). In a variety of natural measures, including typical RT series, the frequency *f* and power *S(f)* are proportional:

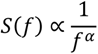

To estimate α, both the frequency and power are log-transformed, and a linear regression line is fit in this log-log space. The linear slope indicates the α value. The power spectrum density was calculated on the entire RT series, using the inverse of the trial number as frequency (following Wagenmakers et al., 2004).

While white noise with an SD of 1 shows a flat power spectrum centred on zero, this is not the case for higher SDs. Instead, when shuffling RT data or when generating random data series with the same variance (Perquin et al., 2020; see https://osf.io/a6zsv/ for simulations), the intercept is dependent upon the overall variance in the series – which is particularly problematic when looking at intra- and inter-individual correlations. To correct for these differences, each RT series was randomly shuffled 100 times. The mean power spectrum of these 100 iterations was subtracted from the original RT power spectrum. Next, the linear regression slope was calculated in log-log space – with the absolute value of the slope representing the exponent α in 1/f^α^. In contrast, the AC1 on the shuffled RT series did behave like white noise (centred around zero).

#### Detrended Fluctuations Analysis

While PSD is meant for “stationary time series – time series that have a constant mean and variance throughout (e.g., white noise) – DFA is supposed to be more robust against non-stationarity (e.g., brown noise; Stadnitski, 2012). Indeed, sensorimotor data series may often be non-stationary – for example, participants may be faster in the latter part of the experiment due to practice effects, meaning that their mean RT is not constant throughout. DFA has gained popularity in cognitive neuroscience in the recent years (e.g., Irrmischer, van der Wal & Linkenkaer-Hansen, 2018; Simola et al., 2017).

To estimate DFA, the time series *x* of total length *N* is integrated into *y(k)* by calculating the cumulative sum of each observation *n* relative to the mean of the time series *μ*:

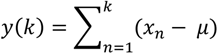

Next, *y(k)* is divided into *b* number of windows *y_b_(k)*. Each *y_b_(k)* value is detrended by the linear trend of that window. Note that if the window sizes are logarithmically spaced, this puts less emphasis on the shorter time scales compared to PSD. On the detrended values, the root mean square error – also called ‘average fluctuation’ *F* – can be calculated as a function of *b* with:

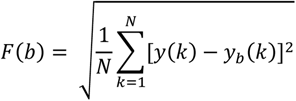

Both *F(b)* and *b* are log-transformed and linearly fit. Regression slope *α* is interpreted as amount of temporal dependency.

DFA was performed on each RT series with the *Fractal* package (Constantine & Percival, 2017), following the procedure of Stadnitski (2012), over non-overlapping blocks log-linearly from a minimum of 4 trials (as lower window sizes are not recommended for linear detrending; Peng et al., 1994) to 512 trials (maximum window size we were able to use). The linear regression slope was calculated in log-log space. Similarly, for each RT series, DFA was performed on 100 randomly shuffled series. The slope of the original RT series was corrected by the difference between the mean slope of the shuffled series and white noise (.5), because the uncorrected values were clearly above chance level for all participants (median value = .57, ranged .56-.58)., though we note they were not correlated with the overall variance of the series. In some papers, the lower frequencies (for PSD) and smallest windows (for DFA) have been excluded – we will address these analysis choices in the section *Control analyses*.

#### ARFIMA models

Although PSD and DFA have been popular for analysing RT, the methods have an important limitation: they have difficulties differentiating long- from short-term dependencies. The use of autoregressive fractionally integrated moving-average (ARFIMA) models has been suggested (Wagenmakers et al., 2004; Torre et al., 2007) as an alternative to statistically *test* the benefit of a long-term parameter—and, as detailed above—ARFIMA allow for predictions which the other dependency measures do not allow. The ARFIMA model is an extension of the ARIMA model, which consists of a combination of three processes, as detailed below.

The first process of the model is the autoregressive process (AR), which aims to capture short-term dependencies. The model takes on an ‘order’ *p*, reflecting how many AR parameters are being estimated. For an AR model of order *p,* AR(p), observation *n* in time series *x* is predicted by its preceding observations *x_n-1_* to *x_n-p_*, with φ_1_ to φ_p_ reflecting the weight for each observation. The model also includes an independently drawn error term ε_n_. As such, the model can be described as:

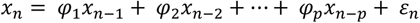

The second process refers to the moving-average process (MA), which also captures short-term dependencies. For a MA model of order *q,* MA(q), observation *n* in time series *x* is predicted by a combination of random error ε_n_ and the error terms of the preceding observations, ε_n-1_ to ε_n-q_, with θ_1_ to θ_q_ reflecting the weight for each error term:

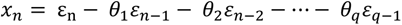

These two processes can be combined into a mixed ARMA(p,q) model:

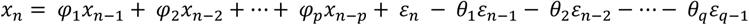

For example, an ARMA(1,1) model can be described as:

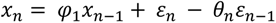

AR, MA, and ARMA are all meant for stationary time series. If a time series is not stationary, an ARIMA(p,d,q) model should be used instead. This model includes a long-term process *d*, referring to the number of times a time series should be ‘differenced’ to make it (approximately) stationary. In the process of differencing, each observation in the time series is subtracted from its subsequent observation. For instance, an ARIMA(1,1,1) model then takes the form of:

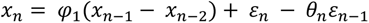

Importantly, in an ARIMA model, *d* refers to a discrete value. Most typical values for d are 1 or 2, which can remove respectively linear and quadratic trends. Instead, in the ARFIMA model the series is instead ‘fractionally differenced’ – such that *d* can take on any value between −.5 and .5. Similarly, *d* in the ARFIMA model refers to a long-term process. One advantage of ARMA/ARFIMA is that they are nested models – which means that the best model can be selected using goodness-of-fit measures such as the Akaike Information Criterion (AIC; Akaike, 1974) and/or Bayesian Information Criterion (BIC; Schwarz, 1978). As such, one can fit both ARMA and ARFIMA on a time series, and test if the long-term parameter *d* sufficiently adds new information (Wagenmakers et al., 2004; Torre et al., 2007).

An ARFIMA(1,d,1) model was fitted on each of the RT series using the *Fracdiff* package (Fraley, Leisch, Maecler, Reisen & Lemonte, 2012), following the procedure of Wagenmakers et al. (2004), to extract the long-term parameter d, together with the weight of the two short-term (first-order) parameters AR and MA (referred to as φ_1_ and θ_1_ in Introduction). These parameters as well as the fit measures (AIC/BIC) are stable across iterations (i.e., when running the pipeline twice, one would get the same values). On the shuffled series, the distributions of d, AR, and MA were all higher than chance (median values respectively .03, .31, and .35), and for the d parameter, these values were weakly correlated with the overall variance of the series. Hence, all three parameters were corrected by subtracting the value of the mean parameters of the shuffled series. The corrected value of AR from five participants and MA from four participants exceeded the theoretical limit (−1) by a small amount (maximum values −1.06 and −1.08). Because we are interested in individual differences, we did not fix these to −1, as to reduce the variation between individuals. To keep the inter-measure correlations as fair as possible, the same 100 shuffled RT series were used for the PSD, DFA, and ARFIMA corrections.

### Archival datasets

The archival MRT data from Anderson et al. (2021) includes a single session of the same task plus an attention-deficit related traits questionnaire on a larger sample. The data from Jin et al. (2019) uses another paradigm traditional in the mind wandering literature (the sustained attention task - SART), in conjunction with a Visual Search, very common in the visual and cognitive literatures. Both tasks include thought probes on attentional state, which allows us to examine their relationship with temporal structure in tasks other than the MRT. Finally, the data from Hedge et al. (2018) are particularly suited for studying reliability, as it includes multiple tasks that all have two sessions. Wagenmakers et al. (2004) is a key paper in the literature on temporal structures, and we show the temporal structures in RT series on each of the six tasks as comparison for our current estimates. Details on methods and data preparation for all datasets can be found in the Appendix.

### Bayesian analyses

Bayesian statistics were conducted in JASP (JASP Team, 2017), using equal prior probabilities for each model and 10000 Monte Carlo simulation iterations. We report the Bayes Factor (BF), which indicates the ratio of the likelihood of the data under the alternative hypothesis (e.g., the presence of a correlation) compared to the null-hypothesis. For example, a BF_10_ of 3 in a correlation analysis means that the likelihood for the data is 3 times larger under the correlation/alternative hypothesis than under the null-hypothesis, and would start to be interpreted as evidence in favour of a correlation. On the other hand, a BF_10_ of 0.33 means that the data is 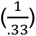 three times more likely under the null than under the correlation hypothesis, which would start to be interpreted as evidence for the null hypothesis. This can also be written as BF_01_ = 3, representing the inverse of BF_10_. A BF_10_ between 0.33 and 3 is typically referred to as ‘indeterminate’. For interpretation purposes, we report BF_10_ when the alternative hypothesis is favoured and report BF_01_ when the null hypothesis is favoured across our main analyses.

In these analyses, the magnitude of the BF refers to the evidence regarding a presence/absence of a correlation. To also get an estimate of the precision of the correlation coefficient, we report the 95% credible intervals (CI) of the posterior distribution alongside our main analyses, which reflects that there is a 95% probability that the correlation coefficient is in said interval.

## Data availability

Our own raw data from the reported analyses is available at https://osf.io/pez34/, alongside the analysis code and jasp files. Any of our other reported measures are available upon request.

## Part 1: Establishing the presence of temporal dependencies

Before examining the intra- and inter-individual correlates of temporal structures in RT series, we first tested whether these series unambiguously showed any temporal structures. Figure 3 shows the distributions of the six temporal dependency measures (AC1, PSD, DFA, and three ARFIMA(1,d,1) parameters) for each task separately. Distributions with the same colour indicate that these tasks were performed by the same group of participants.

**Figure 3.**
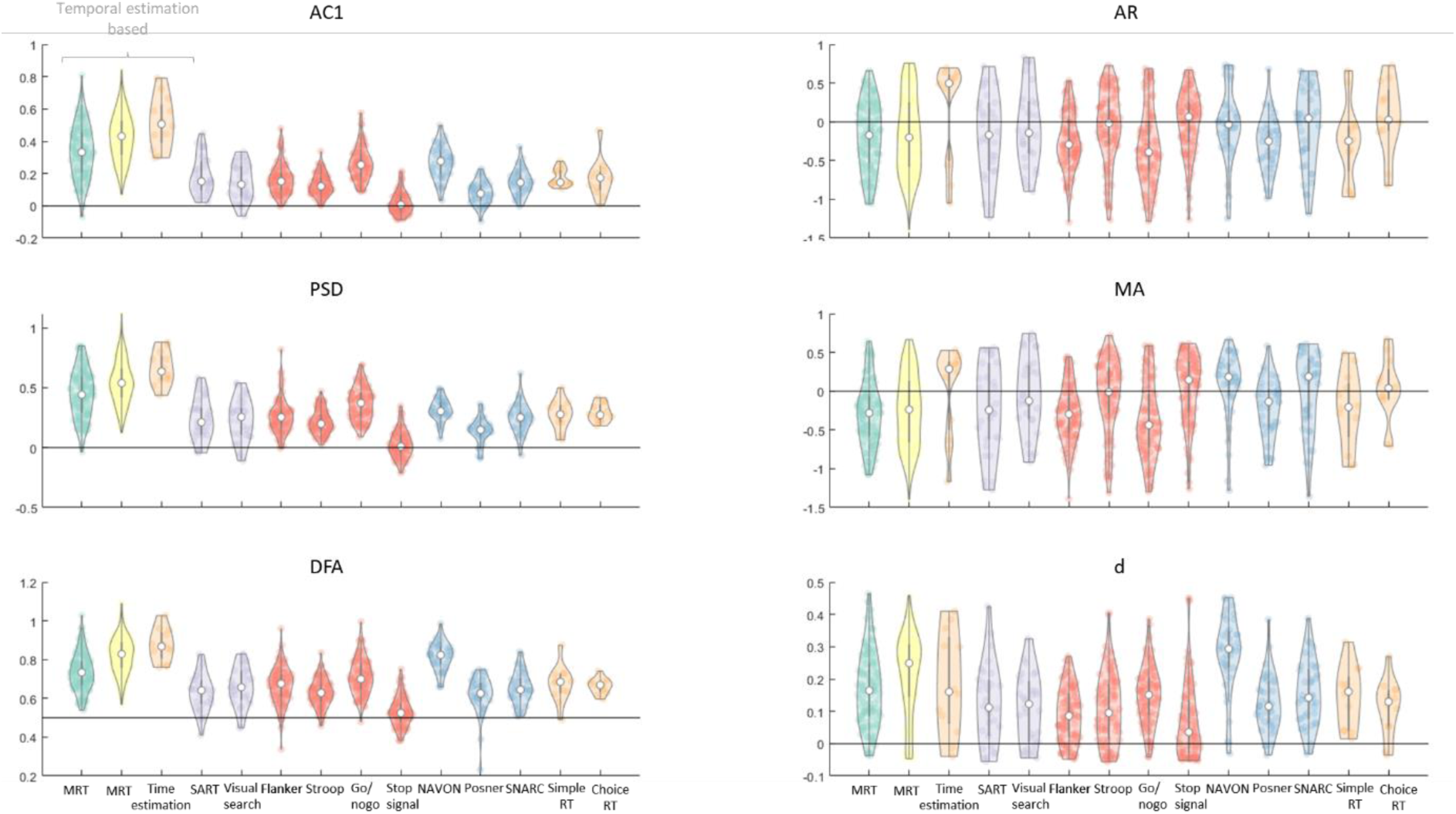
Overview of the temporal structure measures (AC1, PSD slope, DFA slope, and the three ARFIMA(1,d,1) parameters – AR, MA, and d) across all participants of each dataset. Each dot represents a measure from one participant, with the group median of each distribution in white. In each subplot, the horizontal line reflects the null-hypothesis of the featured temporal dependency measure. Colours represent the different datasets: Participants from the MRT data are shown in green (our current data, both cohorts combined, from the first MRT session only) and yellow (from Anderson et al., 2021), participants who did both the SART and Visual Search task (Jin et al., 2019) are shown in purple. For the cognitive tasks, participants who performed the Eriksen Flanker, Stroop, Go/No-Go, and Stop-signal tasks are shown in red, while participants who performed the NAVON, SNARC, and Posner cuing tasks are shown in purple. For comparison, participants on the simple RT, choice RT, and time estimation tasks from Wagenmakers et al. (2004) were also added. We conclude that the AC1, PSD, DFA, and d reflect a clear presence of temporal structure and that, consistent with previous literature, the timing-based tasks show the strongest dependency.

In each subplot, the null-hypothesis (i.e., absence of temporal dependency) is reflected by the horizontal line. For AC1, PSD, AR, MA, and d, this corresponds to 0, and for DFA, the null-hypothesis is .5. Bayesian One Sample t-tests were conducted on AC1 and PSD slopes – to test if they were statistically different from zero – and DFA slopes – to test if they were statistically different from 0.5. On our MRT data and on the cognitive tasks, this was done separately for the first and the second session. We found extreme evidence for the existence of temporal structures for all tasks except the Stop-Signal task in both sessions (Table 1). We performed the same analysis on the ARFIMA(1,d,1) parameters and found extreme evidence for each distribution of d values being higher from zero (see below for formal comparison between ARFIMA and ARMA model). The AR and MA parameters were less consistent across datasets. At first glace, the different measures mostly seem similar to each other, despite their different interpretations, with exception of the AR and MA, which one would expect to resemble AC1. We run inter-measure correlation analyses as sanity check (see section *Control analyses)* and return to the interpretation of the AR and MA parameters in the Discussion. For now, we conclude that there is clear temporal structure in our collected and archival datasets.

Wagenmakers et al. (2004) found that the temporal estimation task elicited stronger dependency than the simple and choice RT tasks. Consistent with this, we find that data the MRT, which is also time-estimation-based, showed relatively high structure.

**Table 1.** Bayes’ Factors in favour of the existence of temporal structures in the RT in the different measures: AC1, the linear fitted slope of the spectral power, the linear fitted slope on the detrended fluctuation analysis, and the ARFIMA(1,d,1) parameters (AR, MA, and d). Green and red fonts indicate clear evidence against and for the null respectively. Black font indicates indeterminate evidence.

|  | AC1* | Spectral slope* | DFA slope** | AR* | MA* | d* |
| --- | --- | --- | --- | --- | --- | --- |
| **MRT data from both cohorts** |  |  |  |  |  |  |
| Session 1 | > 1000 | > 1000 | > 1000 | > 1000 | > 1000 | > 1000 |
| Session 2 | > 1000 | > 1000 | > 1000 | 26 | > 1000 | > 1000 |
| **Data from Anderson et al., 2021** |  |  |  |  |  |  |
|  | > 1000 | > 1000 | > 1000 | > 1000 | > 1000 | > 1000 |
| **Data from Jin et al., 2019** |  |  |  |  |  |  |
| SART | > 1000 | > 1000 | > 1000 | .91 | 1.33 | > 1000 |
| Visual search | > 1000 | > 1000 | > 1000 | .22 | .23 | > 1000 |
| **Data from Hedge et al., 2018** |  |  |  |  |  |  |
| Flanker S1 | > 1000 | > 1000 | > 1000 | > 1000 | > 1000 | > 1000 |
| S2 | > 1000 | > 1000 | > 1000 | > 1000 | > 1000 | > 1000 |
| Stroop S1 | > 1000 | > 1000 | > 1000 | .24 | .23 | > 1000 |
| S2 | > 1000 | > 1000 | > 1000 | > 1000 | > 1000 | > 1000 |
| Go/No-go S1 | > 1000 | > 1000 | > 1000 | > 1000 | > 1000 | > 1000 |
| S2 | > 1000 | > 1000 | > 1000 | > 1000 | > 1000 | > 1000 |
| Stop-Signal S1 | 3.18 | 2.11 | > 1000 | .12 | .19 | > 1000 |
| S2 | 7.87 | .33 | > 1000 | 1.27 | .30 | > 1000 |
| NAVON S1 | > 1000 | > 1000 | > 1000 | .24 | .30 | > 1000 |
| S2 | > 1000 | > 1000 | > 1000 | .24 | 6.44 | > 1000 |
| SNARC S1 | > 1000 | > 1000 | > 1000 | .20 | .17 | > 1000 |
| S2 | > 1000 | > 1000 | > 1000 | .20 | .18 | > 1000 |
| Posner S1 | > 1000 | > 1000 | > 1000 | > 1000 | 40 | > 1000 |
| S2 | > 1000 | > 1000 | > 1000 | > 1000 | 63 | > 1000 |

\* $BF_{\text{value}=0}$ , \*\* $BF_{\text{value}=.5}$

### Long-term dependencies

One benefit of the ARFIMA(1,d,1) model is its direct way to test the benefit of a long-term parameter over only short-term parameters. To test this, the difference in Akaike information criterion (AIC) between the ARMA(1,1), which does not include long-term dependencies, and ARFIMA(1,d,1), which does include long-term dependencies, models was calculated (following the procedure of Wagenmakers et al., 2004). Figure 4 shows this difference for each participant of our MRT data, ordered according to the value of their d parameter. Values above 0 indicate a better (lower) AIC for the ARFIMA model, while values below 0 indicate a better AIC for the ARMA model. In practice, however, only differences larger than 2 are taken as clear support for one model over the other (blue area in Figure 5; Wagenmakers et al., 2004).

**Figure 4.**
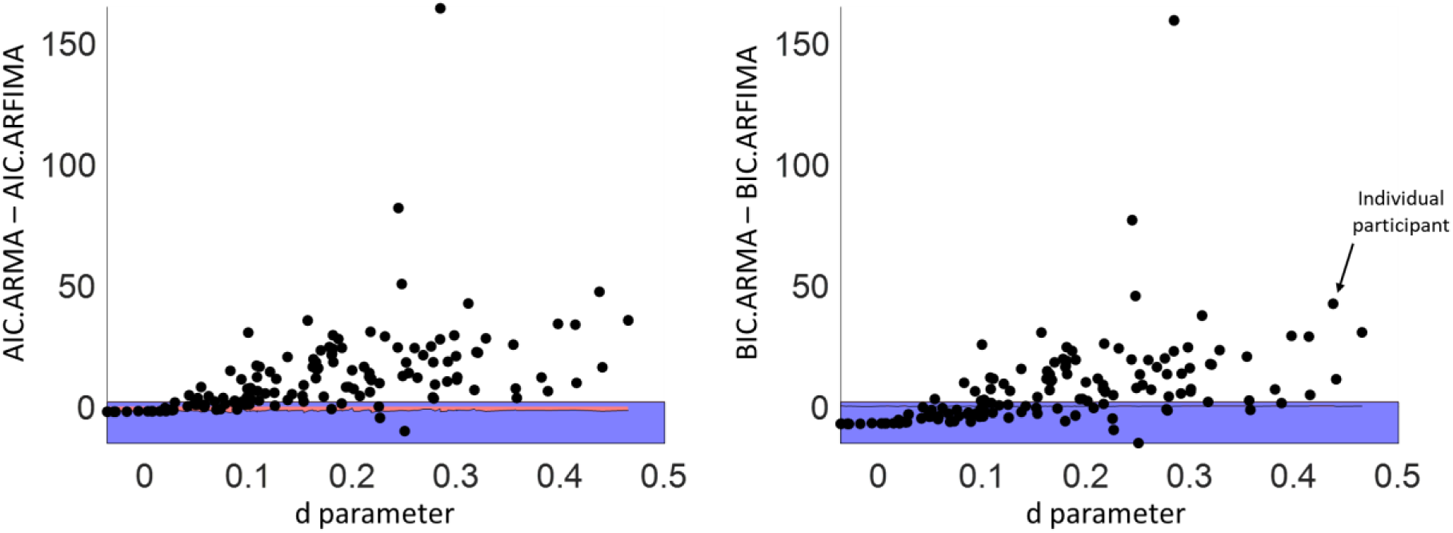
Difference in AIC (left) and BIC (right) between the ARMA(1,1) and ARFIMA(1,d,1) models for the MRT data, with each dot representing one individual subject. For points above the blue-shaded (difference score of 2) and red-shaded (difference score on the shuffled data series) area, the AIC/BIC clearly favours the ARFIMA(1,d,1) model – indicating that the d parameter adds substantial explanation to the model. This was true for most of the participants.

**Figure 5.**
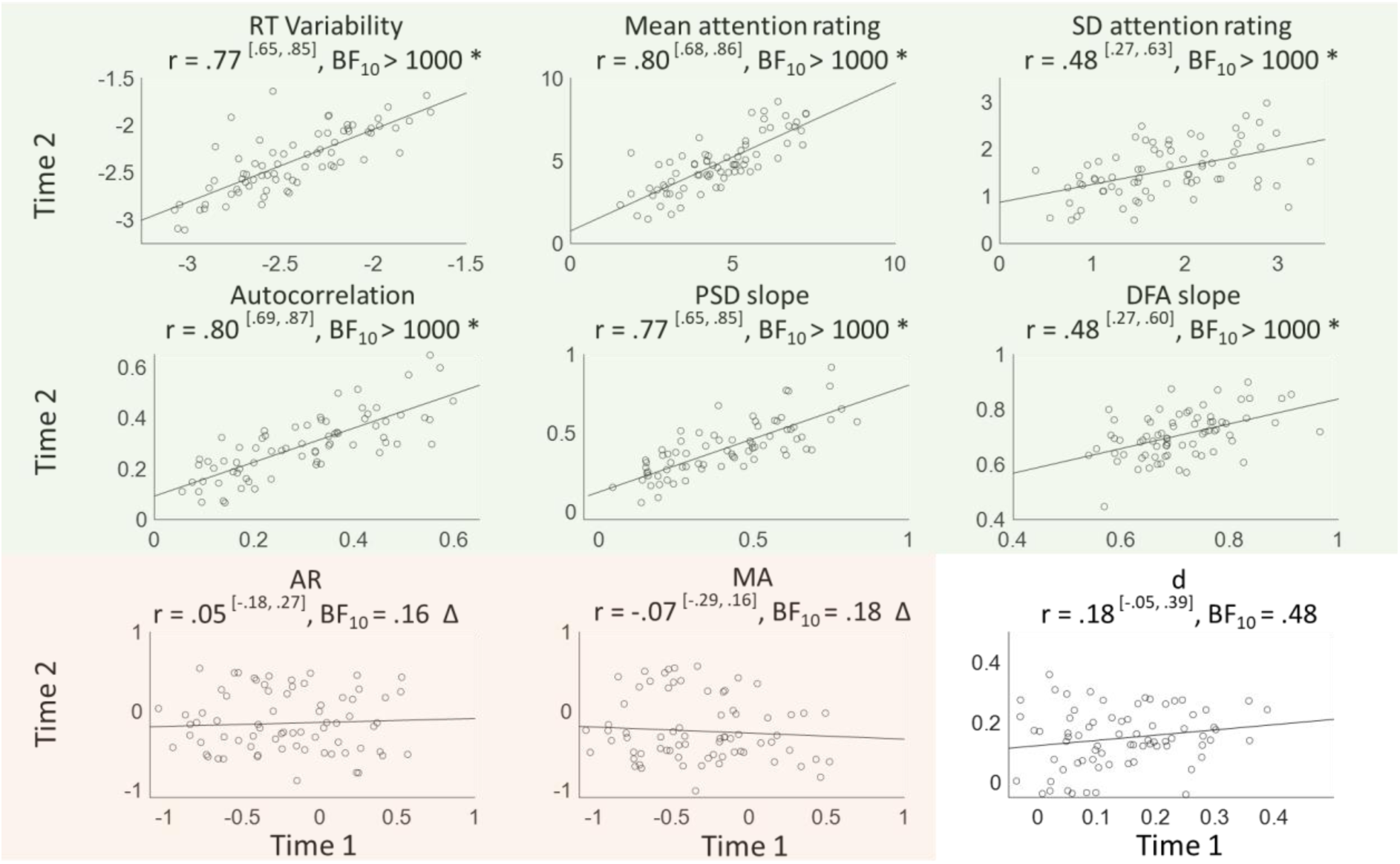
Within-subject correlations between MRT sessions 1 and 2 for performance (RT variability, logged), subjective attentional state ratings (mean and variability), and temporal dependency measures (AC1, PSD slope, DFA slope, and the three ARFIMA(1,d,1) parameters – AR, MA, and d). Values within superscripted brackets indicate 95% credible intervals. Corresponding Bayes Factors above 3 are shaded in green and marked with an asterisk * (indicating clear evidence in favour of that correlation), while Bayes Factors below 0.3 are shaded in red and marked with a triangle Δ (indicating clear evidence against that correlation). For illustrative purposes, PSD slope has been multiplied by −1, meaning that for all temporal dependency measures, higher values indicate more dependency.

Amongst the first MRT session of the 139 analysed participants, the long-term model was clearly favoured for 102 of them (~73%). When using the Bayesian information criterion (BIC) instead, a more conservative goodness-of-fit measure (as recommended by Torre et al., 2007), the long-term model was still clearly favoured for 76 participants (~55%; right panel). The same analyses performed on shuffled RT series show no clear preference for either model for all participants (red area on Figure 5). In the MRT series from Anderson et al. (2021) and the Visual Search and SART RT series, we find similar patterns (Table 3). We conclude that in these tasks, long-term correlations were likely presents in a majority of participants, but still note this was not the case for all of them. Below we assess the stability of these individual differences and explore their potential relevance.

In contrast, for the cognitive tasks, the long-term model was only favoured for the Flanker, Stroop, Go/No-go, and Posner tasks when using the AIC. When using the BIC, the (more parsimonious) short-term model was favoured for all tasks (Table 3). As the one-sided t-tests (Table 1) showed, the temporal dependency measures were clearly different from their null-hypothesis, we can still assess the repeatability and generalisation of these measures. We return to their conceptual value in the Discussion.

**Table 2.**
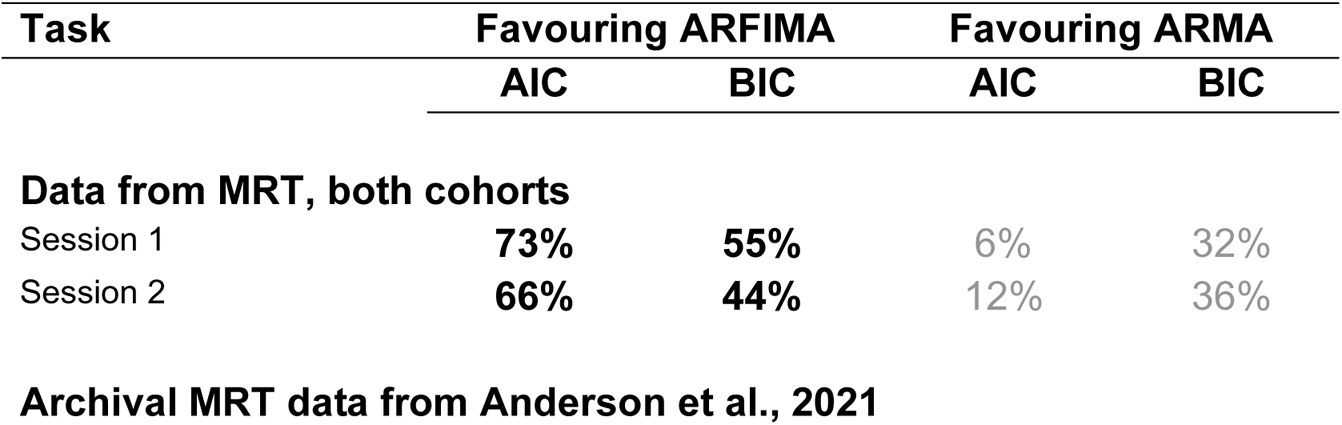
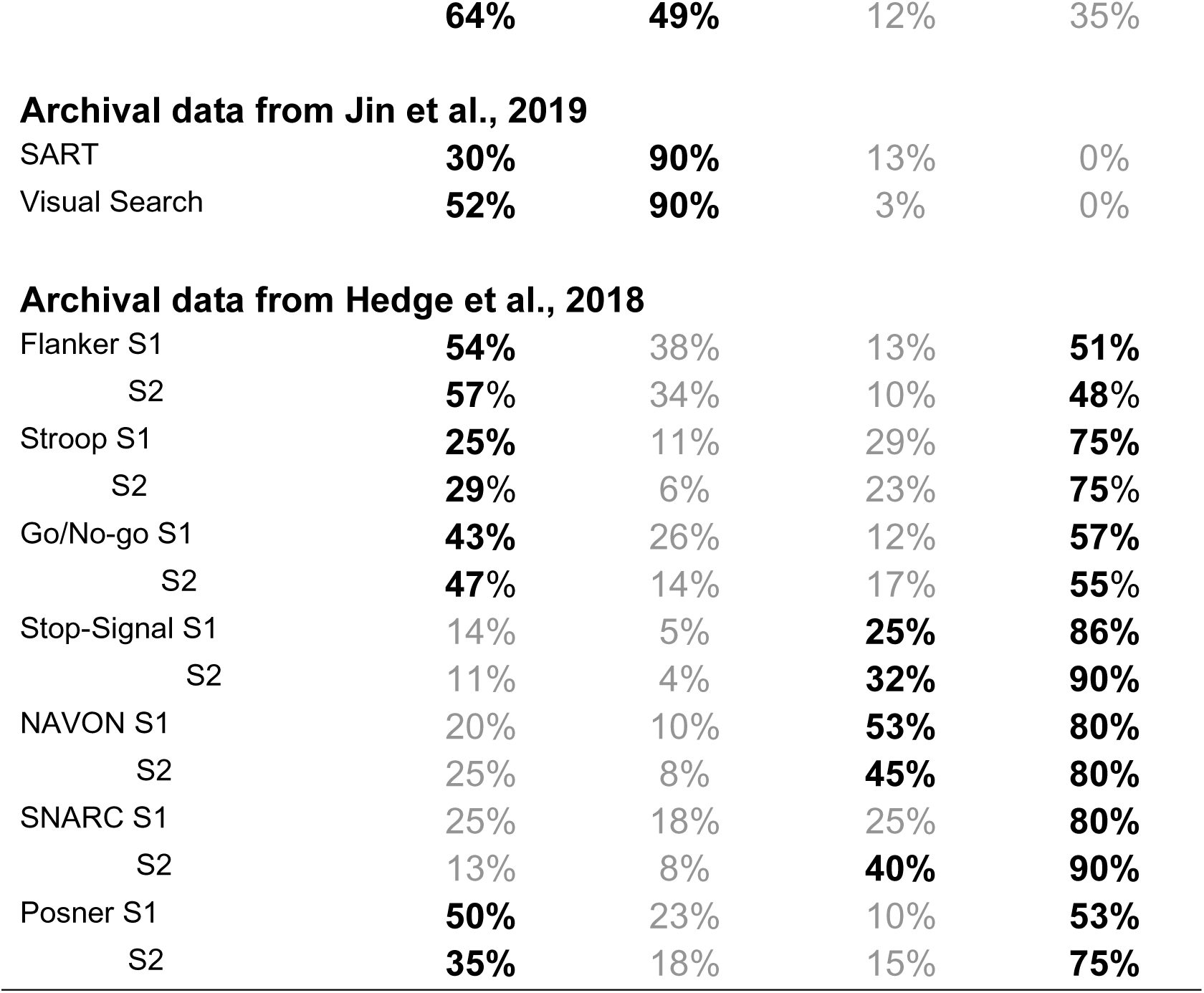
Percentages of participants for whom the fit values clearly favoured the ARFIMA(1,d,1) model (as indicated by a difference of 2 or higher) and for whom the fit values clearly favoured the ARMA(1,1) model (as indicated by a difference of −2 or lower) – separately for the AIC and BIC. For each dataset and each session, the favoured model is shown in bold, and the non-favoured model is shown in grey. S1/S2 indicate the session numbers.

| Task | Favouring ARFIMA |  | Favouring ARMA |  |
| --- | --- | --- | --- | --- |
|  | AIC | BIC | AIC | BIC |
| **Data from MRT, both cohorts** |  |  |  |  |
| Session 1 | **73%** | **55%** | 6% | 32% |
| Session 2 | **66%** | **44%** | 12% | 36% |
|  | **64%** | **49%** | 12% | 35% |
| **Archival data from Jin et al., 2019** |  |  |  |  |
| SART | **30%** | **90%** | 13% | 0% |
| Visual Search | **52%** | **90%** | 3% | 0% |
| **Archival data from Hedge et al., 2018** |  |  |  |  |
| Flanker S1 | **54%** | 38% | 13% | **51%** |
| S2 | **57%** | 34% | 10% | **48%** |
| Stroop S1 | **25%** | 11% | 29% | **75%** |
| S2 | **29%** | 6% | 23% | **75%** |
| Go/No-go S1 | **43%** | 26% | 12% | **57%** |
| S2 | **47%** | 14% | 17% | **55%** |
| Stop-Signal S1 | 14% | 5% | **25%** | **86%** |
| S2 | 11% | 4% | **32%** | **90%** |
| NAVON S1 | 20% | 10% | **53%** | **80%** |
| S2 | 25% | 8% | **45%** | **80%** |
| SNARC S1 | 25% | 18% | 25% | **80%** |
| S2 | 13% | 8% | **40%** | **90%** |
| Posner S1 | **50%** | 23% | 10% | **53%** |
| S2 | **35%** | 18% | 15% | **75%** |

## Part II: Repeatability of temporal dependencies

### Test-retest repeatability in the MRT

Having established that there is substantial temporal dependence in the different tasks we tested, we then sought to determine how stable these temporal dependencies were. To test the intra-individual repeatability of our MRT measures (RT variability, mean and SD of subjective attentional state, and temporal dependency of the RT series), Bayesian Pearson correlation pairs were computed for each measure between time one and two (Figure 5).

Overall, performance (as measured by RT variability over the entire session) and subjective attentional state ratings (mean and variability) showed high repeatability over time. Looking at the temporal structure measures, AC1 and PSD were the most repeatable (equally high as the performance measures). The ARFIMA parameters were unreliable, while the DFA slope fell in-between with the correlation between the two sessions indicating modest repeatability.

### Test-retest repeatability in cognitive tasks

To test the intra-individual repeatability of temporal dependency of the RT series from two sessions in seven well-known cognitive tasks (with 104 participants in the Flanker, Stroop, Go/No-go, and Signal task, and 40 participants in the NAVON, SNARC, and Posner cuing task), Bayesian analyses of Pearson correlations were conducted separately for each measure between time one and two. Figure 6 (left panels) shows the distribution of within-task correlation coefficients (top-left) across all tasks and all measures, with the corresponding BF-values (bottom-left), with each task denoted by a different symbol and the median denoted by the white dot.

**Figure 6.**
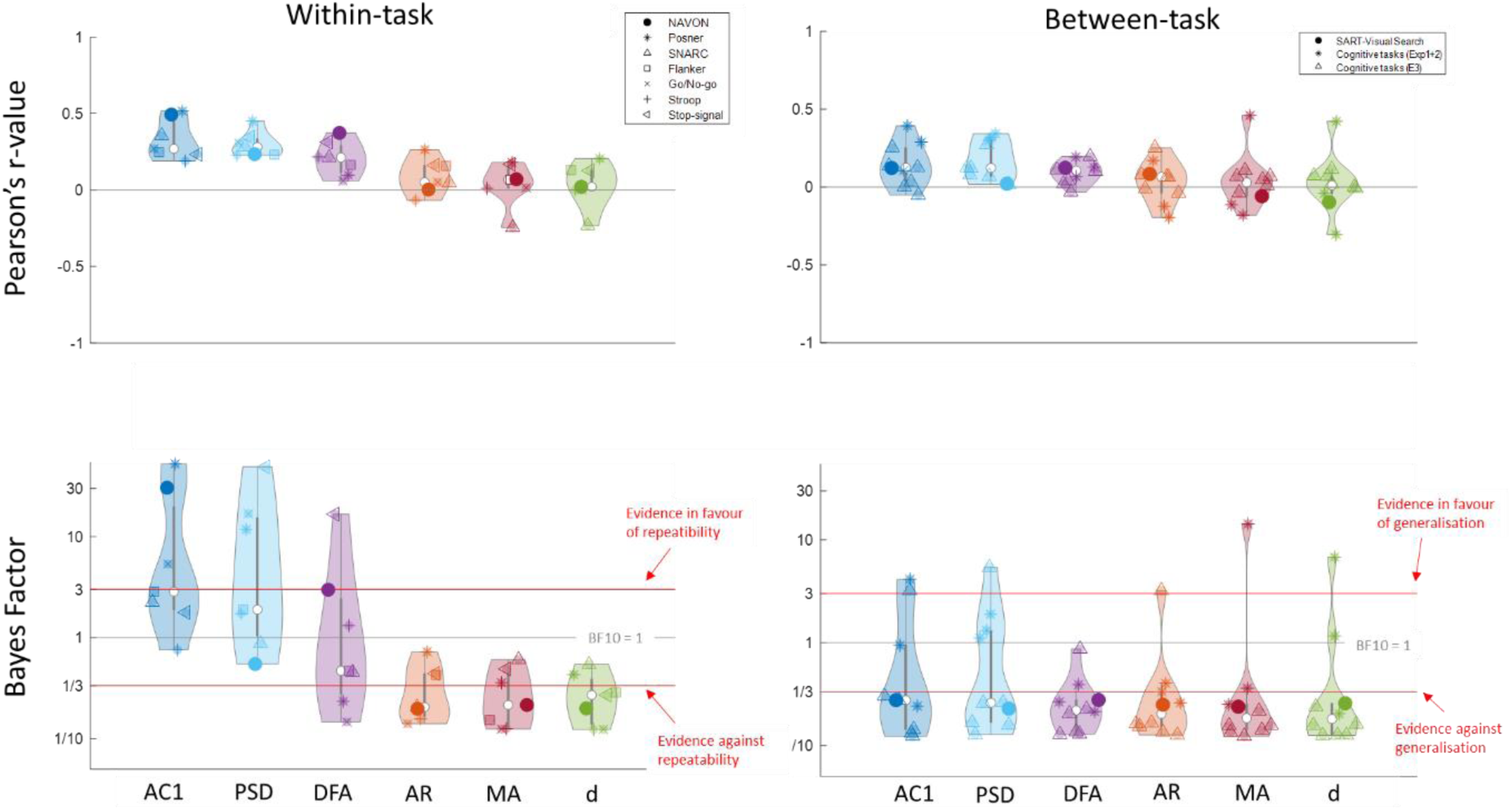
Distributions of correlation coefficients for the within-task (left) and across-task (right) repeatability for the six temporal dependency measures. In each distribution, the white dot represents the median value. The within-task distributions show the correlations between session 1 and session 2 (upper panel) across 104 participants in seven cognitive tasks, with the corresponding Bayes Factors in the lower panel. These reflect stability. The between-subject distributions show the correlations between each combination of tasks for the four cognitive tasks from Exp.1 and 2 and the three cognitive tasks from Experiment 3 from Hedge et al., (2018) as well as from the SART and Visual Search task from Jin et al. (2019). These reflect generalisability. We found that on average, the AC1 and PSD slope were moderately repeatable, while the DFA and ARFIMA parameters were not repeatable. None of the measures were generalisable across tasks. The precise r-values, BF, and 95% credible intervals can be found in the Supplementary Materials.

Similar to our MRT data, AC1 and PSD are the most repeatable of the temporal dependency measures, while the ARFIMA(1,d,1) parameters are clearly not repeatable. However, the repeatability of AC1 and PSD was much lower compared to the MRT, and not consistent across tasks (with only RT series of the NAVON and Posner Cuing task showing moderate repeatability. The DFA slope was not repeatable overall.

### Generalisation of temporal dependencies across tasks

To test the intra-individual generalisation of temporal dependency in RT series between different tasks, Pearson correlation coefficients and corresponding Bayes Factors were calculated for each pair of tasks using the first session of the Flanker, Stroop, Stop-signal, and Go/No-go tasks (104 participants), the first session of the NAVON, SNARC, and Posner cuing task (40 participants), and SART and Visual Search data (30 participants) – all three denoted by a different symbol (Figure 6, right panels).

Out of 60 BF-values relating to temporal dependency, only 5 indicated clear evidence for a correlation, with low to moderate r-values (.25-.43): the AC1 and MA parameter between the NAVON and SNARC, the AC1 and PSD between the Go/No-Go and Stroop, and the d parameter between Flanker and Stroop task. Overall, Bayesian evidence clearly favoured the absence of repeatability across tasks for all measures (Figure 6; bottom-right, with corresponding correlation coefficients shown top-right).

## Part III: Between-subject correlates of temporal dependencies

### Task performance

We then asked whether the various measures of temporal correlation were correlated with task performance. Pearson correlation coefficients and Bayes Factors were calculated between each temporal dependency measure and RT variability on our MRT data (Figure 7, left). We found that participants who performed well (low SD) on the task displayed on average low temporal dependency (as indicated by strong correlations with AC1, PSD and DFA slopes). Crucially, these correlations cannot depend on the variance of the time series, as we correct for this by subtracting the shuffled data series. In the absence of correlation with the d parameter, we cannot conclude that the relationships between RT variability and temporal dependencies are carried by long-range correlations.

**Figure 7.**
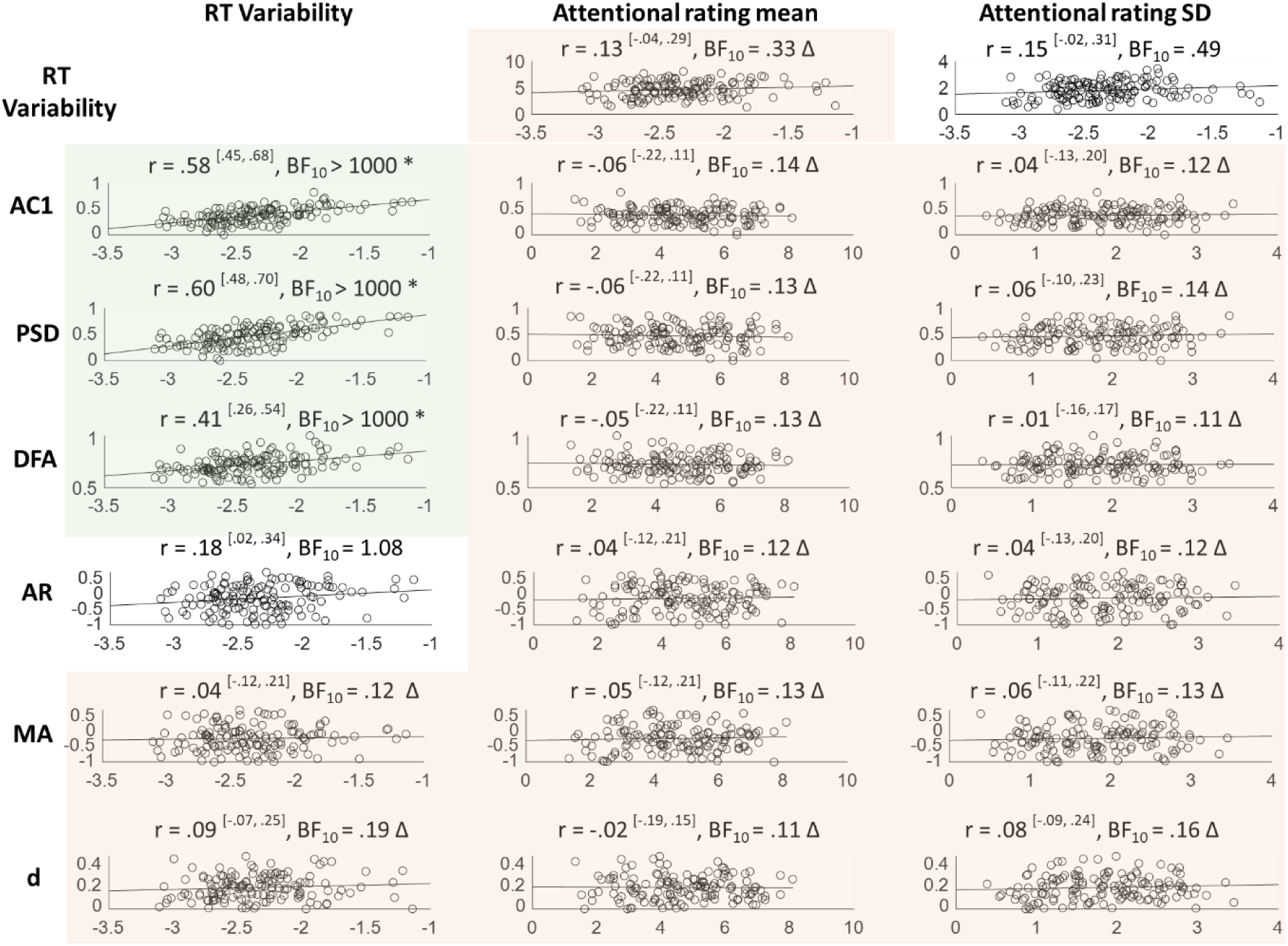
Between-subject correlations between temporal dependency and MRT measures. Values within superscripted brackets indicate 95% credible intervals. Corresponding Bayes Factors above 3 are shaded in green and accompanied by an asterisk * (indicating clear evidence in favour of that correlation), while Bayes Factors below 0.3 are shaded in red and accompanied by a triangle Δ (indicating clear evidence against that correlation). RT variability correlates moderately to strongly with the three repeatable temporal dependency measures (AC1, PSD, and DFA), but not with the ARFIMA(1,d,1) parameters. Bayes Factors show evidence against correlations between subjective attentional state ratings and temporal dependency.

Figure 8 shows these dynamics in more detail for four example participants. Good performance (left column), as indicated by low variability, was associated with a low AC1, that appears to quickly decay over the increasing lags, as well as with relatively shallow PSD and DFA slopes (note that DFA slopes for white noise are .5). Poor performance on the other hand (right column), as indicated by a high SD, was associated with high AC1, that appears to decay only slowly over the next lags, as well as with relatively steep PSD and DFA slopes. Average performance (respectively showing SD around median and mean values) showed intermediate temporal structures.

**Figure 8.**
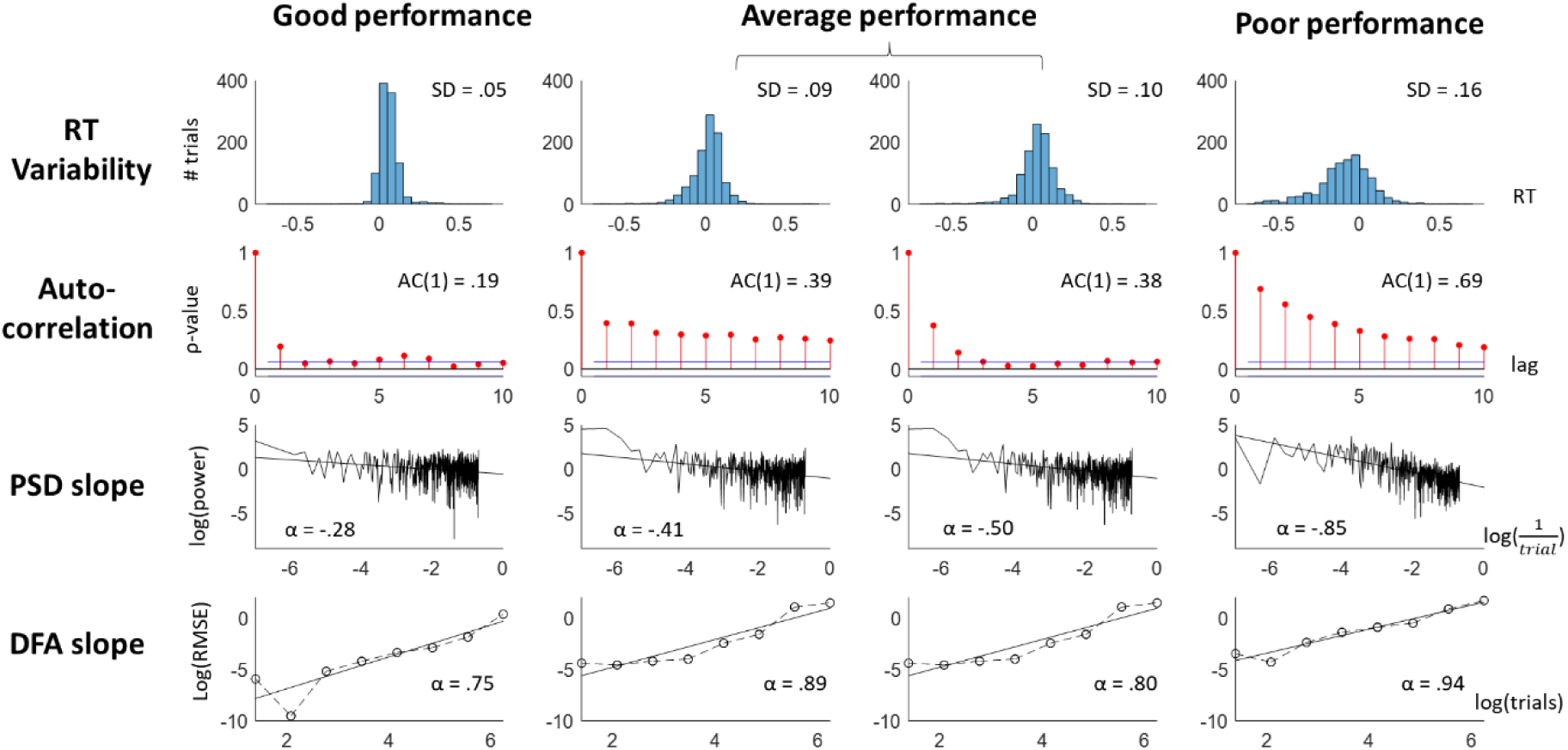
Examples from four participants with (from left to right) good, close-to-median, close-to-mean, and poor performance. Shown for each participant are (from top to bottom) their RT distribution, autocorrelations at the first 10 lags, Power Spectral Density with fitted slope, and Detrended Fluctuations with fitted slopes.

To verify if our results were not confounded by strategy (e.g., trying to anticipate the tone versus responding to the tone), we first reran our analyses after excluding those participants who had a mean RT below −100 ms or above 100 ms (28 participants in total), leaving only the participants who are good at the task. This approach gave highly similar results. Next, we created four subgroups based on low/high variability (<45TH and >55th percentile of the group distribution) and high/low autocorrelation (similar rule). Each subgroup showed the same overall patterns. Our results thus held up well across different strategies.

#### Archival MRT data

As discussed in the *Data preparation and Analysis* section, the RT variability from the Anderson et al. (2021) is not a pure measure of performance, but rather mixes performance and strategy. For consistency, we also computed Bayesian Pearson correlation analyses between the temporal dependency measures and the RT variability on these MRT data, but these should be interpreted with caution.

The correlation patterns in these MRT series were mixed: the AC1 was not correlated to SD, the PSD showed a weak positive correlation, and the DFA showed a weak negative correlation (Figure 9). Reflecting strategy, mean RT was also considered, and showed overall negative correlations with temporal dependency measures (r = −.31, −.38, −.11; BF_10_ > 1000, > 1000, and .53 for AC1, PSD, and DFA respectively) – indicating that participants who were less instruction-compliant (i.e., higher RT) had on average less temporal structure. Again, RT variability was not correlated with any of the ARFIMA(1,d,1) parameters (Figure 9).

**Figure 9.**
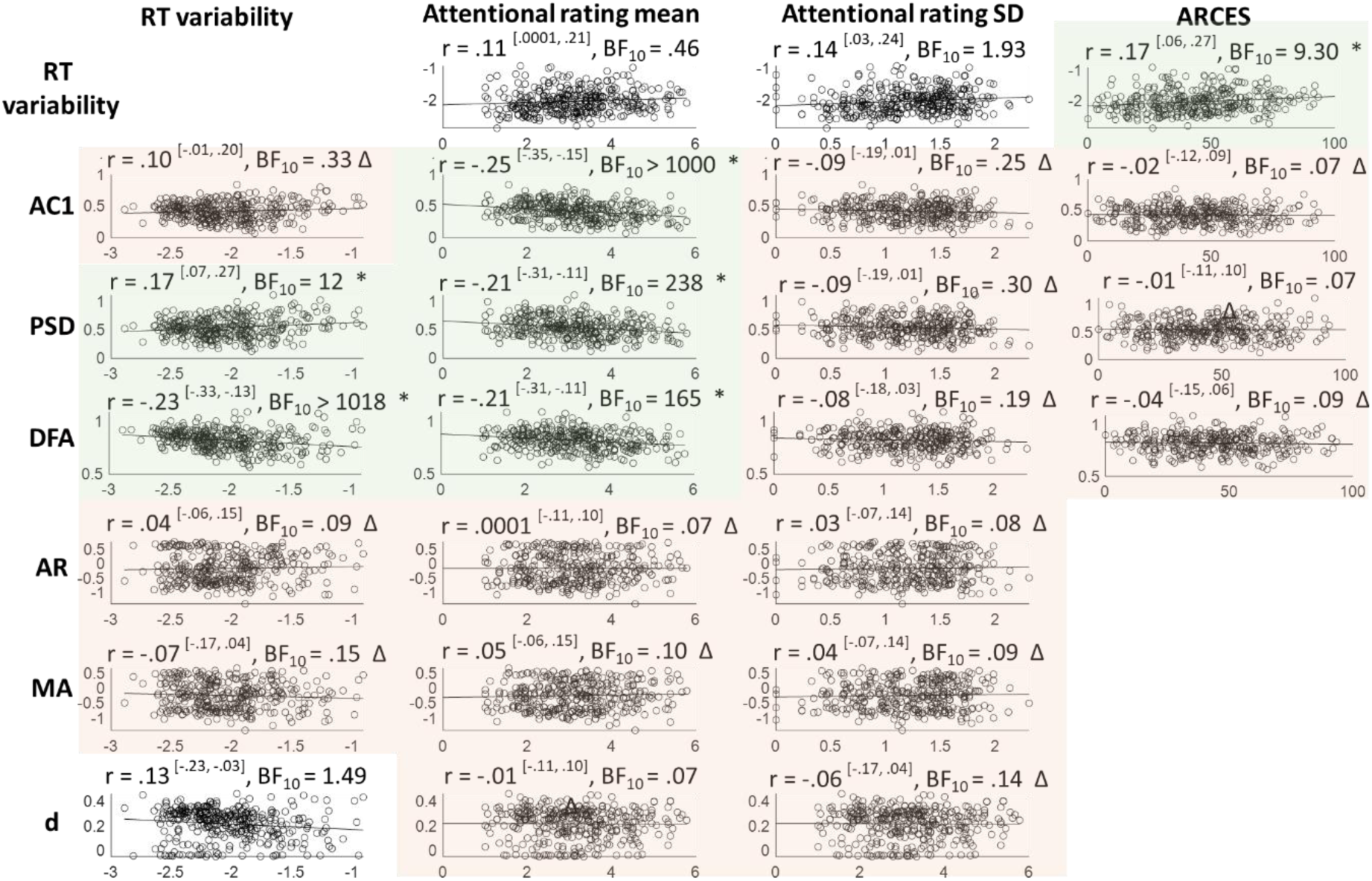
Between-subject correlations of temporal dependency with objective and subjective MRT measures and the ARCES scores, using the MRT data from Anderson et al. (2021). Values within superscripted brackets indicate 95% credible intervals. Corresponding Bayes Factors above 3 are shaded in green and accompanied by an asterisk * (indicating clear evidence in favour of that correlation), while Bayes Factors below 0.3 are shaded in red and accompanied by a triangle Δ (indicating clear evidence against that correlation). Mean subjective attention correlated negatively with AC1, PSD, and DFA. The self-assessed attention-deficit traits correlated positively with RT variability, but not with the temporal dependency measures. ARFIMA parameters were not included in external validity analyses, as they were not repeatable.

### Subjective attentional state

Bayesian Pearson correlation analyses were conducted between each temporal dependency measure and the mean and SD of the attentional state ratings on our MRT data (Figure 7, right), and consistently indicated evidence against any correlations. For comparison, the correlation between RT variability and subjective ratings was also included.

#### Archival MRT data

When running the same correlation analyses on the archival MRT data, there was clear evidence that mean attentional state ratings correlated negatively with the AC1, PSD, and DFA measures (Figure 9, middle two columns) – indicating that higher reports of being off-task were associated with less temporal structure – but not with the ARFIMA(1,d,1) parameters. Numerically, this correlation was strongest for AC1, though all three coefficients were in the low range. Neither AC1, PSD, nor DFA was correlated with the SD of attentional state ratings.

#### Visual Search and SART

The same correlations were run between the proportion of off-task probes and each temporal dependency measure separately for the SART and the Visual Search data. There was evidence against any correlations (BF_10_ ranging .22-.46 for the SART, with 5 out of 6 showing determinate evidence; BF_10_ ranging .23-.54 for the Visual Search, also with 5 out of 6 showing determinate evidence).

### Self-assessed attention-deficit related traits

After examining correlations with objective and subjective task-based attention measures, we then asked whether the temporal dependencies correlated with more global self-reported attentional traits. To this end, Bayesian Pearson correlation analyses were conducted between self-assessed attention-deficit related traits and the repeatable temporal dependency measures (AC1, PSD, and DFA). For comparison, RT variability was again included. Neither self-assessed ADHD traits (both cohorts) nor daydreaming and impulsivity in daily life (cohort 1) correlated with RT variability (Figure 10).

**Figure 10.**
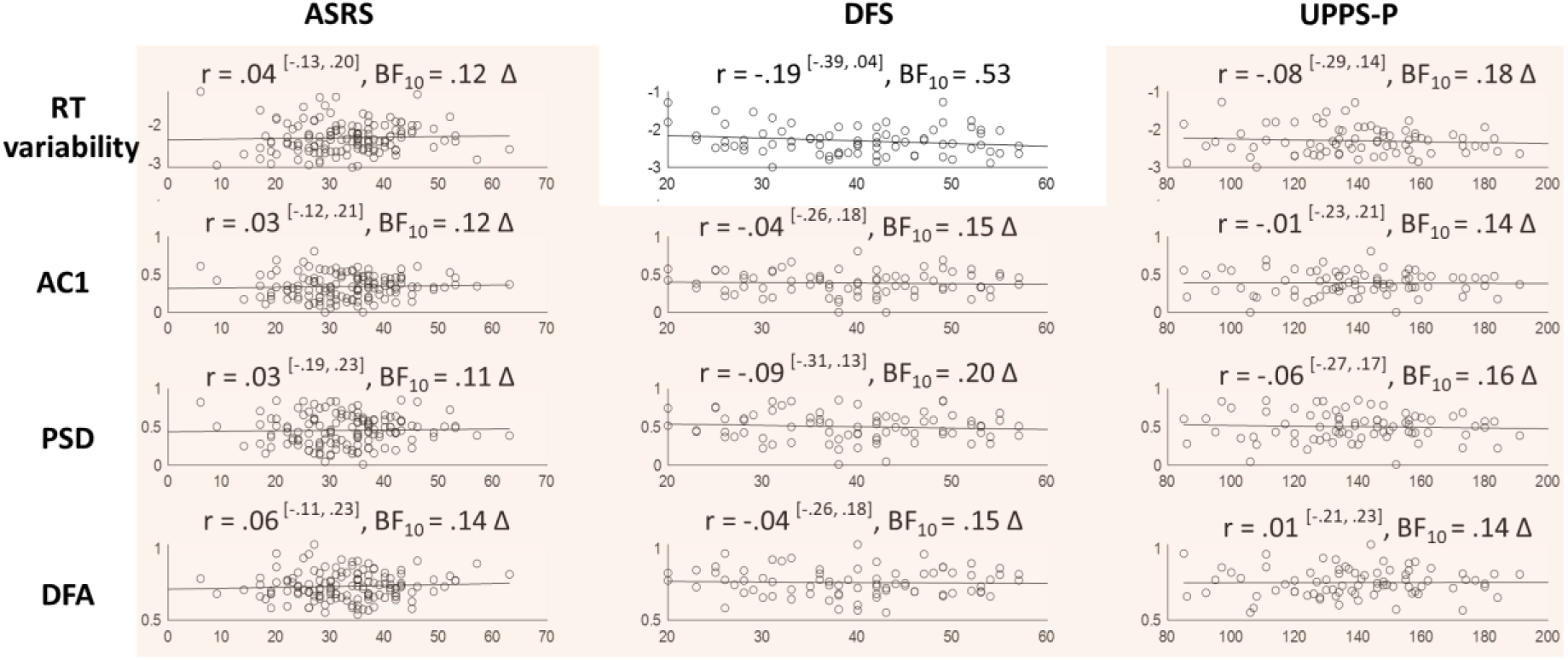
Between-subject correlations between the repeatable temporal dependency measures (AC1, PSD, and DFA) and self-assessed attention-deficit related traits. Corresponding Bayes Factors above 3 are shaded in green and accompanied by an asterisk * (indicating clear evidence in favour of that correlation), while Bayes Factors below 0.3 are shaded in red and accompanied by a triangle Δ (indicating clear evidence against that correlation). Eleven out of twelve pairs showed clear absence of correlation (one indeterminate).

#### Archival MRT data

The self-assessed scores of attention-related cognitive errors in daily life modestly correlated with RT variability (Figure 9). However, the scores did not correlate to any of the repeatable (AC1, PSD, DFA) temporal dependency measures.

## Control analyses

### Conceptual check: Inter-measure correlations

As we were interested to see how well the different measures of temporal dependency correlated to each other, Bayesian Pearson correlations across the different temporal dependency measures of our MRT data (performed on the data from session 1) revealed that the measures that showed some within-individual repeatability (AC1, PSD slope, and DFA slope) strongly correlated with each other (Table 4). Counterintuitively maybe, although the AR and MA parameters from the ARFIMA model were not repeatable, they still correlated highly with each other (but not to any of the other measures, including the conceptually similar AC1).

**Table 4.** Pearson correlations with 95% credible intervals within superscripted brackets, and corresponding BF_10_ between the different measures of temporal dependency on our MRT data. Green and red fonts indicate clear evidence against and for the null respectively.

|  | PSD | DFA | AR | MA | d |
| --- | --- | --- | --- | --- | --- |
| **AC1** | .96 [.94, .97]<br>(1.3<sup>e+72</sup>) | .82 [.75, .86]<br>(2.2<sup>e+31</sup>) | .16 [-.01, .32]<br>(.63) | -.01 [-.18, .15]<br>(.11) | .38 [.22, .51]<br>(3139) |
| **PSD slope** | - | .74 [.65, .81]<br>(2.9<sup>e+22</sup>) | .18 [.01, .33]<br>(.90) | -.04 [-.20, .13]<br>(.12) | .26 [.10, .41]<br>(14.4) |
| **DFA slope** | - | - | -.04 [-.20, .13]<br>(.12) | -.06 [-.22, .11]<br>(.14) | .62 [.50, .71]<br>(2.2<sup>e+13</sup>) |
| **AR** | - | - | - | .93 [.90, .95]<br>(3.3<sup>e+57</sup>) | -.19 [-.34, -.03]<br>(1.32) |
| **MA** | - | - | - | - | .04 [-.13, .20]<br>(.12) |

The high correlations of AC1 with PSD and DFA and that between AR and MA were also consistently present in the archival datasets (Supplementary Materials B). The d parameter showed more volatile correlations with the other measures.

### Analysis choices

Here we discuss how the current results patterns hold up with different analysis choices. It is impossible to validate the current against all possible analysis choices, so we limited ourselves to two factors that we deemed most important (see Discussion): the choice of frequency or length in the long-range estimates, and the method to deal with missing data and/or outliers.

Firstly, we calculated PSD over all the possible frequencies. However, log-log spectra of behaviour tend not to be linear all the way through, but instead increase in power at the highest frequencies – resulting in a small curve at the high end of the spectrum (see Torre & Delignières, 2008 Figure 1, for instance). Some previous studies therefore exclude the highest frequencies, to fit the slope only on the linear part of the spectrum. In line with Torre et al. (2019), we excluded 1/8^th^ of the highest frequencies and reran the within- and between-subject analyses. Similarly, we recalculated the DFA measures with a minimum window of 16 trials (e.g., 16-512 trials instead of 4-512).

Secondly, the current results are based on the time series with the missing values excluded. For the MRT data, we reran the analyses with two different methods: 1) replacing the missing values with each individual’s median RT, and 2) replacing with an RT of 650 ms (reflecting the maximum time a participant had to respond). For the cognitive, SART, and Visual Search tasks, we only reran the analyses with a median imputation, as there is no obvious ‘extreme value’ in any of these tasks.

Overall, the patterns were fairly robust to the different analysis choices, though least so when the missing values were replaced with the highest possible value (.650, for the MRT data only). The full results are shown in Supplementary Materials C, in which all noteworthy exceptions (i.e., with correlation coefficients being at least .10 higher or lower than the original value, or correlation coefficients of which the corresponding Bayes Factor switched our evidence categories) are highlighted.

## General Discussion

We provide the first large-scale investigation of the repeatability and inter-individual correlates of temporal structures in behavioural time series. To do this, we contrast the most commonly used methods, applied to rich multi-measures data that allow us to conjunctly assess, on the same participants, performance and their temporal structure, subjective attentional state, test-retest repeatability and personality traits. The take-home message, based on our own data, is illustrated in Figure 11, where repeatable variables are shown in bold, and the proximity between two variables captures the strength of their correlation.

**Figure 11.**
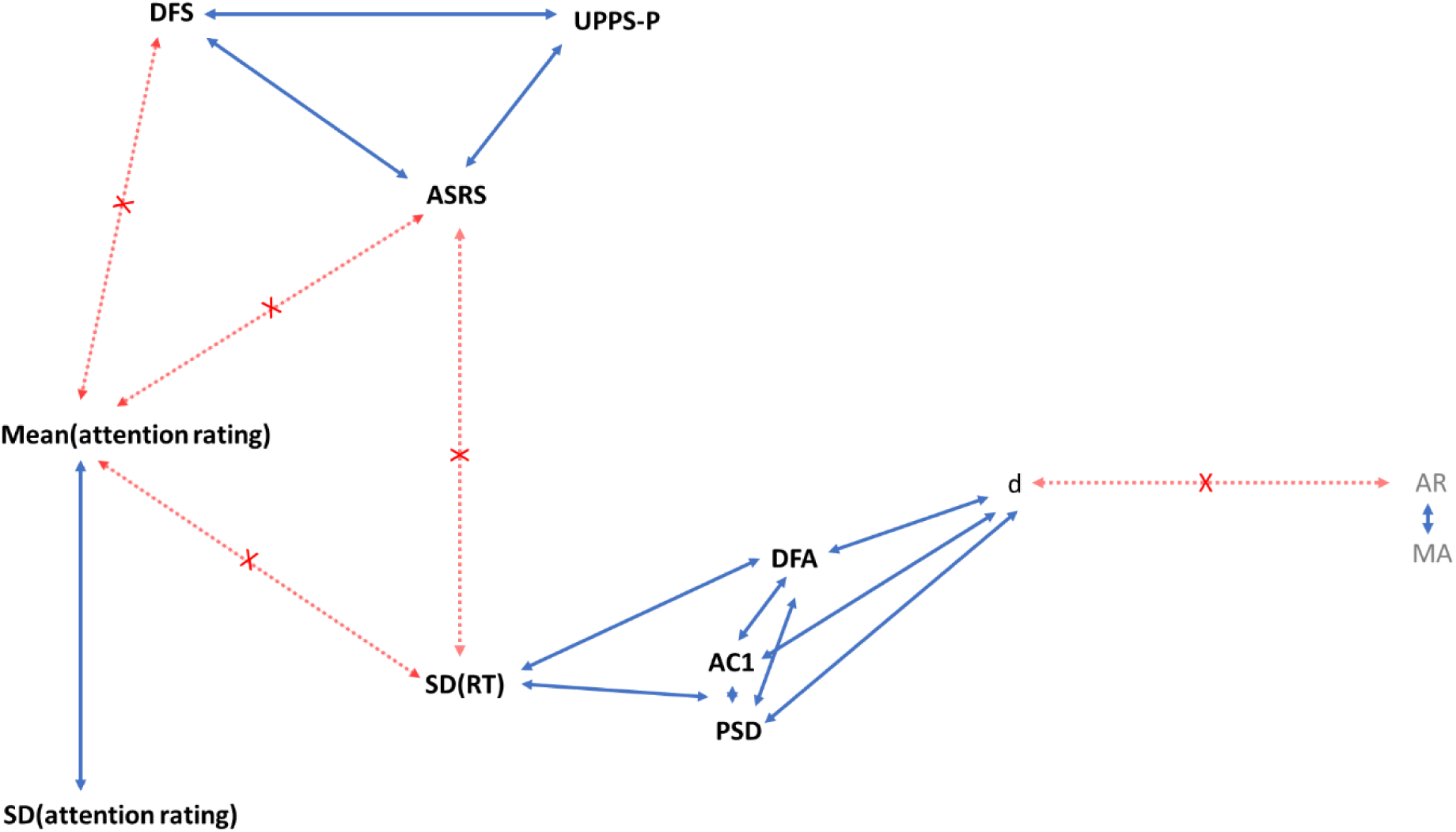
Illustration of the repeatability and relationships across all the measures from our MRT data. Measures written in bold show at good intra-individual repeatability (or moderate for DFA), as found either by present between-session correlations, or from the literature for the questionnaires. Positively correlated variables are linked by a blue arrow with a length proportional to 1 – r, with r being the Pearson’s r correlation coefficient (see Figures 3–7, 9–10), i.e., strongly correlated variables are shown close together. The most notable absences of correlations are flagged with a red line and cross. These may have been expected based on literature – such as the relationship between ADHD tendencies and RT variability – or on the supposed similarity between the measures – such as within-task attentional state ratings and general mind wandering tendencies (DFS). Note that none of the archival datasets considered offered a broad enough range of measures to justify a similar figure.

We found that temporal dependency showed repeatability over time, though this was dependent on which measure was used. AC1 and PSD were highly repeatable, and were the only measures that showed the same strength of correlation as the objective measure of behavioural variability and the subjective measure of mean attentional state. Instead, the DFA slope was only moderately repeatable, and the ARFIMA parameters not at all. The temporal dependency measures (with exception of the ARFIMA(1,d,1) parameters) did correlate with performance – such that good performance was associated with less temporal structures. However, there was Bayesian evidence against correlations of the temporal dependency measures with both subjective attentional state and self-assessed personality traits. In Figure 11, this is reflected by ‘cluster forming’ of the different variables: A cluster of questionnaire-measures, a cluster of attentional state ratings, and a cluster of behavioural RT and its features – indicating poor external validity.

To investigate the generalisability of our results, we further considered three archival datasets. While these were designed for very different purposes, each of them could contribute to a subset of our research aims. These showed that temporal structures are observable in most other cognitive tasks, though with a substantially lower magnitude compared to the MRT, that they are overall repeatable but not generalisable across tasks, nor correlated with self-assessed attention-deficit traits.

### Intra-individual repeatability

Our conclusions that DFA slopes show moderate within-task repeatability at best and does not translate across paradigms is consistent with previous work (Smit et al., 2013; Torre et al., 2011). Computing a Cronbach’s α on our MRT data between the two sessions gives a value (α = .47) in the same range as Torre et al.’s. Here we extend their results by including other measures of temporal dependency. Our results indicate that AC1 and PSD measures were more repeatable (with respectively ~64 and 59% of shared variance across sessions, while the DFA slopes only share ~22% of variance), but also did not translate across paradigms. By extending this conclusion to more reliable measures, we can make a stronger case for the lack of generalisation.

These repeatability scores are comparable to those reported in the neural domain for temporal structures of alpha and beta power in resting-state EEG recordings (split-half reliability: Smit et al., 2013; test-retest reliability: Nikulin & Brismar, 2004). Interestingly, individual variations in these measures seem partially driven by genetic factors, as shown in two adolescent twin studies which estimated the heritability of DFA slopes (Linkenkaer-Hansen et al., 2006; Smit & Anokhin, 2016). To what extent these heritability estimates extend to temporal structure in behavioural data remains an open question.

### Temporal dependencies, performance, and attentional state

Temporal dependency (as indicated by higher AC1, PSD, and DFA) increased with poorer performance. Superficially, this appears in line with Smit et al. (2013) and Irrmischer et al. (2018) and at odds with Simola et al. (2017). One way to bring all results in line with each other is by assuming that the temporal dependencies reflect mental flexibility, rather than task performance per se. We can speculate that participants with low mental flexibility perform better on the Metronome Task because they stick to the consistent action throughout – resulting in a negative between-subject correlation between good task performance and temporal structure. In turn, participants with low mental flexibility would be perform poorly on the Go/No-Go task because they are bad at switching between responses – resulting in a positive between-subject correlation between good task performance and temporal structure.

In practice however, it can be difficult to distinguish which tasks require ‘flexibility’ and which require ‘consistency’, as performing most psychological tasks requires a careful balance of different – often clashing – task demands (e.g., we want to be as fast as possible, but also not make any mistakes). For example, in the Go/No-Go task from Simola et al. (2017), participants are required to make a response on 75% of the trials (Go-trials) and to abstain from responding on all other trials – and good performance may therefore rely heavily on response inhibition. In the Continuous Temporal Expectations Task from Irrmischer et al. (2018), participants also only respond to Go-trials, but these targets were rare, appearing only every fourth to tenth trial – and good performance may therefore rely heavily on sustained attention. Still, either task clearly requires some of both elements, and it is difficult to see how the different task demands would lead to this particular pattern of results.

Findings across different tasks may also be difficult to compare because switches between different experimental conditions across trials might affect the measurement of temporal dependency. This could affect different tasks to different extents (or not at all in tasks such as tapping which feature no experimental condition). To check for these effects, we corrected RT series for differences in conditional mean RTs in the Flanker and Stroop tasks, as well as the NAVON, SNARC, and Posner tasks, by subtracting the difference between the condition mean and the grand mean from each RT (e.g., if the grand mean = 400 ms, the mean of condition A = 410 ms, and the mean of condition B = 390 ms, then every RT from condition A would be deduced with 10 ms and every RT from condition B would be increased with 10 ms). Distributions of the temporal dependency measures for original and corrected were virtually the same, and rerunning the repeatability and generalisation analyses resulted in highly similar patterns with only small and unsystematic changes. As such, the measurement of temporal structure does not seem to substantially altered by switches in experimental conditions. We note though that tasks with interleaved conditions do appear to give rise to lower temporal dependencies in general compared to tasks without conditions, as suggested by the distributions of measures across tasks in Figure 3. This seems sensible, as performing a congruent versus incongruent task is not only “doing the same thing but more slowly”, but likely involves different underlying processes (e.g. conflict detection, inhibition), which could very well be less temporally dependent than repeating the same action over and over again.

From a mechanistic perspective (Torre & Delignières, 2008; Torre & Wagenmakers, 2009), one could expect that the relationship between performance and temporal dependencies across participants may strongly depend on the task, and may not be linear. For instance, an engaging task over the course of which one high-performance state dominates would be characterised by low temporal dependencies (because alternations between states are scarce). In this speculative scenario, individuals impacted by more low-performance episodes would see their overall performance decrease but exhibit more temporal structure. In contrast, a task allowing on average for an even split between the two modes may lead to a quadratic relationship, whereby any departure from the average leads to less temporal dependencies.

In the current data, the monotonic increase of temporal structure with RT variability may appear consistent with the following (highly speculative) reasoning: good performance in the MRT task is associated with the dominance of one high-performance mode, while increasingly poorer performance is driven by more switching to a low-performance mode. The *within*-individual local correlation between RT variability and attention ratings suggests that high and low modes could be somehow associated with different subjective feelings of being on task. However, this correlation is weak, and the within-subject distributions of subjective reports were not bimodal. Moreover, we found evidence against correlations between temporal dependency and the *variability* of subjective attention ratings. Therefore, although the idea of mode-switching between on-task and off-task states is seducing, its ability to accurately describe our data is unclear.

### Clinical relevance

To our knowledge, the current study is the first to directly relate temporal dependency to self-assessed personality traits. Bayesian analyses showed evidence against correlations with ADHD tendencies, mind wandering tendencies, and impulsivity. It is important to note that the current study used healthy participants, who do not typically report clinical levels of ADHD symptoms. Oversampling for high ADHD tendencies, testing clinical samples versus healthy controls, or testing the effects of medication on clinical samples may have led to different conclusions.

Gilden & Hancock (2007) have previously compared temporal structures of participants with high versus low RT variability on a mental rotation task. They report that no one in the ‘low’ group reported ADHD symptoms, while participants in the ‘high’ group did. However, a range of methodological issues with the analysis method (see Farrell et al., 2006 for a critique), and the study design (including small sample size, uncontrolled differences between groups, unclear self-constructed questionnaire) make the findings difficult to interpret. Other studies have also looked at the temporal dynamics of RT in ADHD (e.g., Castellanos et al., 2005; Geurts et al., 2008; Johnson et al., 2007; see Karalunas et al., 2012; 2014 for reviews; see Kofler et al., 2013 for a meta-analysis), but their methods and aims have been different than described in the current research. Rather, assuming that people with ADHD are indeed more variable than neurotypicals, these studies have examined whether this increased variability is driven by rhythmic fluctuations – i.e., if the longer RTs observed in ADHD are temporally predictable. Typically, the RT series are transformed to the frequency domain by either a Fast-Fourier or Morlet-wavelet transform to obtain a power spectrum – to test if specific frequencies show higher peaks for ADHD patients compared to controls. Although these studies typically found increased power in ADHD patients, they have only focused on low frequencies (< 1.5 Hz). It remains unknown if this would translate in higher *slopes* (which would mainly happen if lower frequencies were increased but higher frequencies were not). Nowadays, this analysis seems to have gone out of fashion, as differences between groups could not be traced back to one specific low-frequency peak (Karalunas et al., 2012).

In contrast, ADHD in children has been associated with *reduced* autocorrelations in RT compared to healthy samples (Aase & Sagvolden, 2005; Aase, Meyer & Sagvolden, 2006). However, both studies used reinforcement learning tasks – for which performance may exhibit both positive and negative autocorrelations, with the temporal structures reflecting the training process. This makes conclusions from learning tasks hard to generalise to other tasks designed to focus on endogenous variability alone.

Attention deficits have also been investigated in relation to temporal dependency in neural activity (Smit et al., 2016) in adolescents. Attention deficits were measured with a questionnaire that was completed by the mother (The Strengths and Weaknesses of ADHD Symptoms and Normal Behavior Scale; SWAN-AP), and scores were validated with a diagnostic interview. Results indicated no significant correlations between attention problems and DFA slopes in α, β, and θ oscillations in resting-state EEG, though some exploratory analyses suggested the *change* in temporal structure across age may differ for high and low attention-problems groups, with the high group showing more increase of structure over age. Whether these patterns can be extrapolated to adults remains an open question.

The disconnect between task behaviour and self-assessed scores in healthy individuals has been discussed previously, e.g., in the domain of self-control (Enkavi et al., 2019). It has been suggested that this disconnect may be caused by measurement error, as the self-assessments are much more reliable than behaviour. This suggestion cannot be applied to our current findings: both the self-reports and the temporal dependencies were highly repeatable within individuals, and still, they were not informative of each other – suggesting they may measure different constructs altogether.

### Statistical power

As the current paper deals with multiple research questions and a large number of analyses, we used a hard cut-off of 3 for BF in all our figures and interpretations. Most of our BFs were much higher than the cut-off, and we largely find the same patterns across different measures (e.g., different temporal dependency measures, different measures of subjective attentional state, different analysis choices), despite variations in important design factors, including (but not limited to) number of trials, trial length, number of participants, and experiment length – each of which affects the estimates of temporal dependency and the statistical power. Overall, it seems unlikely our findings are due to chance.

The only noteworthy exception relates to correlations between mean subjective attention score and temporal dependency, clearly unsupported in our data but supported in Anderson’s. Although the sample size in Anderson’s data was higher than ours, the BFs in favour of the null-hypothesis in our data were between 7-8 when using the standard priors, providing reasonable support for an absence of correlations. Additional robustness checked showed that the evidence favouring the null-hypothesis was robust to most reasonable prior settings. In other words, it does not appear that our data lack sensitivity, as this would correspond to indeterminate BF_10_-values between 0.33 and 3. Instead, our data are most consistent with the conclusion that there is no effect. It is thus unlikely that the discrepancy results from a lack of power. It rather seems that our participants were performing the task more seriously than Anderson’s, rarely report being off-task intentionally, while this was very common in Anderson’s data. The subjective levels of being off-task from both datasets may thus also be qualitatively different.

### Temporal dependency across the brain-cognition system

Neural oscillatory activity also clearly shows temporal dependencies, as consistently reported in the EEG and MEG literature. However, exact relationships to individual differences in health condition, task performance or temporal structure in behavioural series are difficult to summarise. Linkenkaer-Hansen et al. (2001) presented the first empirical evidence for temporal structure in α oscillations during rest in MEG data, showing the presence of non-zero PSD slopes and of positive autocorrelations that slowly decayed over > 100 s. Typically, the temporal structures appear strongest with eyes closed, and reduce with eyes open and during task (e.g., Irrmischer, Poil, Mansvelder, Sangiuliano Intra & Linkerkaer-Hansen, 2018B; Linkenkaer-Hansen, Nikulin, Palva, Kaila & Ilmoniemi, 2004; Smit et al., 2013), reinforcing the idea that they result from internally-generated dynamics. These neural temporal structures during rest have been studied on their own (without any task to relate to), including in clinical groups such as depression, schizophrenia, epilepsy, Alzheimer’s disease, Parkinson’s disease, and autism (see Zimmern, 2020 for an overview). It is unclear how informative these are to understand individual differences in performance during sensorimotor or cognitive tasks. Below we focus on work that is more directly related to our current focus.

Several studies have reported links across individuals between temporal structure in neural activity and temporal structure in behavioural time series or simply with overall performance, but no consistent picture appears to have emerged yet. One study reported strong between-subject correlations between temporal structure in behavioural hit-miss series during an audio-visual detection task, and temporal structure in MEG oscillatory activity, both at rest and during the task (Palva et al., 2013). In contrast, Smit et al. (2013) reported only weak correlation between temporal structures in finger tapping series and EEG oscillations during rest, and none with EEG activity recorded during the task. Broadening up to individual differences in overall task performance, in the two studies using a sustained attention task, higher temporal structures in resting-state EEG oscillations were associated with better (Irrmischer et al., 2018B) or worse performance (Herzog, Steinfath & Tarrasch, 2021). In contrast, temporal structure of EEG *during the task* was associated with worse performance (Irrmischer et al. 2018B) or not associated with performance (Herzog, Steinfath & Tarrasch, 2021). In conclusion, and similar to the literature on behavioural temporal structure, the literature on neural temporal structures does not currently support clear directional predictions.

### Short-versus long-term dependency

Two overarching questions for both the behavioural and neural temporal structure remain: 1) how persistent are these dependencies over time and, relatedly 2) what are the generative processes driving the presence of dependency? The presence of long-term dependency cannot simply be assessed with DFA and PSD slopes tested against a null-hypothesis assuming no temporal dependency at all (also see Farrell et al., 2005; Wagenmakers et al., 2004), but need to be tested against short-term models.

In the current study, we used short-term autocorrelation and ARFIMA(1,d,1) modelling alongside PSD and DFA. We observe that, even in the tasks in which the fit measures clearly favoured the long-term model (like both MRT tasks), there were plenty of individuals for whom the short-term model was still clearly preferred (ranging from 6-12% of participants for AIC and 32-36% for BIC). Furthermore, in four out of the nineteen group comparisons shown in Table 3, the short-term only model was unambiguously favoured by both AIC and BIC over the model with a long-term parameter for the majority of participants, and in many of the other tasks, support for either model was ambiguous across fit measures. This occurred even when the distributions of group distributions for AC1, PSD and DFA slopes were clearly above their null-hypothesis (Figure 3) – indicating that merely assessing the magnitudes of temporal dependency measures is not sufficient for assessing long-term processes.

Although DFA and PSD measures have been developed to capture temporal dependencies beyond the very-short term correlations captured by the AC1, we observed that all were highly correlated. In datasets where no clear long-term structure was observed, this would suggest that the information driving individual differences in DFA and PSD comes mainly from short-term dependency. This could explain why the correlations between AC1 and the long-term measures are so high.

### Individual differences in subjective attentional state

Subjective ratings of attentional state were highly repeatable across the two MRT sessions, showing ~61% shared variance, but did not appear to transfer between the SART and the Visual Search task (there were no thought probes in the other datasets considered). The self-reported mind wandering in daily life score (as measured by the DFS) has been previously found to be repeatable even after a 1-year interval (Giambra, 1980). However, we found that the attentional state ratings did not correlate with the DFS nor with any other measures across individuals (except for the standard deviation of these ratings). Previous work has reported a similar absence of relationship (Kane, Smeekens, Meier, Welhaf & Philips, 2021) or significant but small correlations (Mrazek et al., 2013; Seli, Risko & Smilek, 2016; Smeekens et al., 2016). This relationship is necessarily expected, as they measure different constructs: the ratings measure the dynamic attentional states throughout a repetitive experimental task, while the questionnaire measures people’s recollection of their overall attention during daily life. Still, as these measures both reflect subjective judgment on attention, a stronger and consistent relationship might be expected. However, it is possible that participants use different criteria for answering the thought probes and questionnaires – for example, one might consider how their attention compares to others when answering questions about their attention in daily life, but consider how their attention compares to their own baseline when answering questions about their attention in the present moment. In that case, weak to no correlations between the two measures would not be surprising. Still, individual differences in subjective attention might be driven by other constructs not measured in the current design.

Rather than the reliability of the subjective ratings themselves, there may be interest in the *relationship* between the subjective ratings and behavioural variability. A strong relationship between behaviour and attentional state ratings may suggest an ability to accurately monitor one’s own internal states. As a control analysis, we quantified the relationship between SD and attentional state for each participant (for details, see Methods) separately for the first and second session, and computed the correlation between them. Results suggested an absence of repeatability (r = .22, BF10 = .87), showing that the relationship of one’s subjective judgments of attention and variability is not a stable trait. Below, we offer two possible (but not necessarily irreconcilable) explanations.

First, within-person stability of subjective ratings may be driven by persistent individual biases in the self-report scales. Research on report biases has mainly been conducted in the context of survey questions, including the social desirability bias, the acquiescence bias (i.e., the tendency to agree with a statement regardless of its content), and the central-tendency bias to use mid-point rather than extreme values. Attempts to overcome these biases have been sought in scale design (e.g., length of scale, verbalisation of categories) as well as Item-Response Theory modelling of the responses (e.g., Kreitchmann, Abad, Ponsoda, Nieto & Morillo, 2019; Menold, 2021; Nadler, Weston & Voyles, 2015; Primi, Hauck-Filho, Valentini & Falk, 2019; Soto & John, 2019). Responses styles and biases differ reliably between individuals (Cronbach, 1946), and may affect the external generalisation of self-reports. To our knowledge though, there is no research particularly on reporting in subjective attentional state in general, and it is unclear how the known biases and their proposed solutions would directly translate to thought probes that are repeatably presented during a task. In research on retrospective subjective confidence of performance, methods have been developed to dissociate ‘metacognitive bias’ (i.e., absolute confidence score) from ‘metacognitive sensitivity’ (i.e., confidence score in comparison to performance, such that an individual with high sensitivity reports high confidence following correct trials and low confidence following errors; Fleming & Lau, 2014). One might expect that reporting biases affect the former but not the latter, but this remains speculative for now, and it likewise remains unclear if the proposed solutions would work for these measures.

Biases may also be introduced by the way that subjective attentional state is measured. Indeed, there is a large variety in probe-based measures (e.g., categorical versus continuous, fewer versus more response options) with little convergence towards a standard (see Weinstein., 2018 for a review). Our measure consisted of a 9-point scale from ‘completely on’ to ‘completely off task’. The benefit of such a scale is that it can measure a larger variability in subjective attention compared to categorical responses, and thus should be able to capture the ‘depth’ of the off-task focus. However, these depth measures appear to be more confounded by the confidence participants have in their ratings compared to measures targeting the content of their thoughts (Kane et al., 2021). Furthermore, the external validity of depth responses has recently been disputed. Ideally, one might expect that any increase in subjective rating is on average associated with worsening of performance. This seems untrue in practice: using a 5-point scale, Kane et al. (2021) only found a significant increase in RT variability between ratings 4 and 5. Similar patterns can be found in our data: most participants do not show a linear relationship between subjective ratings and RT variability.

Secondly, the high repeatability of subjective attentional state and of behavioural variability could be independent of each other. Although the intra-individual relationship between attentional state ratings and variability seems robust, their shared variance is typically very low (including in the current paper, in which the median Kendall’s τ correlation coefficient was .08). Furthermore, while thought probe methods interrupt participants to rate their subjective state, participants are not good at catching themselves being off-task as it occurs (Franklin, Smallwood & Schooler, 2011; Schooler, Reichle & Halpern, 2004), and are unable to use the fluctuations in their attentional state to improve upcoming performance (Perquin et al., 2020). If the intra-individual relationship between subjective attentional state and behavioural variability is weak, it makes sense that any potential reliability of this relationship would be difficult to capture. One approach here may be to examine if the relationship between subjective attentional state and behavioural variability can be increased with interventions. For example, subjective attentional state and behavioural variability have been found to moderately decrease with mindfulness training (Morrison, Goolsarran, Rogers & Jha, 2014; Mrazek, Franklin, Phillips, Baird & Schooler, 2013), but to our knowledge, no study has looked into any changes in the relationship between attentional state and behavioural variability directly. Though the computational methods used to dissociate bias versus sensitivity in confidence ratings do not directly translate to probe-based attentional state ratings (as they are based on accuracy and require a rating for every trial), the attentional state literature may benefit from a similar conceptual approach of testing the relationship between behaviour and subjective ratings directly.

One more general caveat of using thought probes to measure attentional state is that they could themselves affect participants’ RT series and attentional state: each probe itself is a disruption from the actual task. Two studies concluded to the absence of effect of thought probe frequency on objective performance, although they disagreed on the effect on off-task ratings, where one showed that higher frequency reduced the tendency to be off-task (Seli, Carriere, Levene & Smilek, 2013), while the other reported evidence against such effect (Robison, Miller & Unsworth, 2019). Still, these results only pertain to the overall performance, and not to the structures in the time series – it is likely that the temporal structures are affected. The same would be true for pauses between blocks of trials. To our knowledge, pauses have been largely ignored in the literature so far, and more empirical evidence would be needed to assess their effects. In the current study however, participants all received the same number of probes throughout the task, and the lag between probes should on average be equal, meaning that these disruptions to the time series cannot explain our current results in any systematic way. Future studies focusing on individual differences in subjective attentional state may consider keeping the interval between probes consistent between participants to potentially reduce individual variance.

### Different measures of temporal dependency

All of the analyses methods used in the current study are so-called ‘fractal methods’ and are mathematically derivable from each other (Stadnitski, 2012). Similarities in results may therefore be expected. Still, we found important differences over the methods, both in their properties (repeatability and relationship to performance) as well as in the extent to which they correlated with each other.

Our choice of methods was dictated by those previously used on cognitive data. However, these methods: 1) are not exhaustive – other methods, such as Rescaled Range Analysis and Dispersion analysis, fall under the same subclass (see Delignières et al., 2006; Delignières, Torre & Lemoine, 2005 for overviews) – and 2) may come with several variants and refinements. Furthermore, the analyses methods in the current research are all for capturing linear trends in the data over different time windows. Non-linear methods may capture more nuanced temporal trends in the data, and have been used previously on RT data (see Kelly, Heathcote, Heath, & Longstaff, 2001). Again however, these methods have been hardly used on psychological data, and overall, non-linear trends in RT series are difficult to capture, as their presence seems to depend on particular tasks demands and characteristics (e.g., short versus longer inter-stimulus interval; Kelly et al., 2001).

Particularly striking is the extremely high correlation between AC1 and the PSD-slope, with around 92% shared variance. This implies that when studying individual differences, fitting a slope over the power of the *entire* time series (in this case: a range of 1050 trials) gives little additional information to simply correlating each trial to the next. It is clear that the PSD method is not more informative, despite being more computationally heavy, less intuitive in interpretation, and hence more difficult to implement in practical contexts (e.g., physicians working with patients).

Comparisons between the goodness-of-fit of the ARMA(1,1) to the ARFIMA(1,d,1) models (Torre et al., 2007; Wagenmakers et al., 2004) showed the ARFIMA models were favoured on MRT data – indicative of the presence of long-term structure. However, it should be noted that, as the ARFIMA parameters were at best poorly repeatable within individuals, the model may be more difficult to interpret. One possibility for this lack of reliability is that the model estimates three parameters at once. However, the AR and MA weights from the ARMA model (hence only two parameters) were not more repeatable than the AR and MA weights from the ARFIMA model. As a control analysis, we fitted each parameter separately (e.g., fitting an ARFIMA(0,d,0) to estimate *d*) to see if this would improve repeatability. These analyses showed high repeatability across sessions for all three parameters (AR: r = 0.80, BF_10_ > 1000; MA: r = 0.78, BF_10_ > 1000; d: r = .81, BF_10_ > 1000). Furthermore, unlike within the ARMA and ARFIMA model, the single AR weights were the exact same values as AC1 – as one would expect.

As such, the individual parameters get altered when estimated together to obtain a better numerical fit. While this is not necessarily surprising, it does raise questions about the biological plausibility of the model: As short-term dependencies in behaviour (and neural activity) are much easier to explain than long-term dependencies, modelling may instead take an approach in which the short-term parameters are fitted first, and the contribution of a long-term parameter is assessed afterwards.

While AC1 and PSD have the highest repeatability, the DFA may come with most flexibility. One can decide on how many time windows to take into account, and whether these should overlap or not. The fitted slope can be plotted against the window size (see Figure 8 for examples), which allows one to directly assess the fit. Based on this fit, the window size can be adjusted (see Kantelhardt, Koscielny-Bunde, Rego, Havlin & Bunde, 2001; Krzemiński, Kamiński, Marchewka & Bola, 2017 for examples). This ensures the obtained slope actually matches the data – something which is not clear in the PSD slopes (Wagenmakers et al., 2004; Torre et al., 2007). However, this flexibility also has its drawbacks: It opens the door to selective reporting, and can make it more difficult to compare and replicate findings across studies. For example, Irrmischer et al. (2018) used windows of 2-60 RTs on the go-trials (but note that go-trials only occurred every 4 to 10 trials, which means that their DFA slopes are not calculated on the basis of adjacent trials) with 50% overlap between the windows, while Torre et al. (2011) used a maximum window of 256 trials (on a series of 512 trials) without overlap, and Simola et al. (2017) used windows of 30-300 seconds without overlap. While none of these analysis choices are necessarily wrong, it is clearly difficult to compare these findings, which stands in the way of replicability. Ideally, it should thus be reported how any analysis choices were decided upon, and potentially, how different choices may or may not alter the results.

### Missing values in the time series

Regarding the extraction of the different measures, the issue of missing values (i.e., missed responses on trials) has been scarcely addressed. Three methods have been discussed to deal with this issue in our analyses: 1) exclude the missing values entirely from the series (which appears the most common option in the literature), 2) replace the missing values by values that stay true to the distribution of non-missing values (for instance by using the median value, or a value obtained by statistical interpolation; see Adamo et al., 2015 for an example), or 3) replace the missing values by the most extreme value (e.g., the maximum response time).

Overall, the result patterns were fairly robust across these three methods. Still, some substantial changes occurred even when the number of missed responses was low for most participants (group median < 1% in our MRT data). One explanation for these increases in repeatability may be that the number of omissions is itself a repeatable trait (r = .65, BF_10_ > 1000) – although this would not explain why the increase is not found in all the measures. As there is no straightforward way of dealing with these missing values, it may be recommended to also report alternative methods – particularly when the number of missed responses is high and/or different across the groups that are being compared.

The issue of missing values has been mentioned previously by both Kofler et al. (2013) and Karalunas et al. (2012; 2014). They rightly point out that the use of different methods across articles complicates results comparison. We would like to take this one step further: As soon as the time series have a lot of missing values, interpretation becomes more difficult no matter which method is used. This is due to what missed responses possibly represent: extreme cases of poor task performance. By excluding the missing responses or by replacing them with average values, it appears that the participants are doing better than they actually are – by disregarding the moments in which they were doing the task so poorly that they did not respond at all. In other words, imputation of missing values only gives unbiased estimates when the missing values are ‘missing at random’, which is typically not the case in these experimental tasks – which means that there is no reliable way of estimating their values (see Donders, van der Heijden, Stijnen & Moons, 2006 for a review on data imputation). By replacing the missing values with the most extreme values, this issue is solved, as the missing values are being represented by extremely poor performance on that trial. However, this method takes a toll on the RT distributions, and conceptually only works if there is a known maximum (as in the MRT, or in task with a response limit).

It should be emphasised that this problem is not trivial – particularly when studying clinical samples compared to healthy controls. It is a fair expectation that clinical populations show more missing responses – meaning that any method of dealing with the missing values may introduce or mask systematic group differences unrelated to the temporal structures in the time series.

### Our recommendations for studying temporal dependencies

The current study is a large-scale investigation into the repeatability and between-subject attention-related correlates of temporal structures in sensorimotor variability, featuring RT series from 11 tasks (or 19 sessions), using four common temporal structure measures. Based on our findings and experiences with quantifying the dependencies, we make the following six recommendations:

1. *Formally assess the presence of long-term dependency.* The presence of long-term dependency cannot be assumed to be a ubiquitous phenomenon across all individuals and tasks but should be tested against a null-hypothesis that includes a short-term parameters (as opposed to white noise), echoing Wagenmakers et al. (2004; 2012) and Torre et al. (2007).
2. *Formally assess the reliability of temporal structures.* Repeatability of the autocorrelation and PSD slope across time was high in some tasks (time estimation) but lower in other cognitive tasks. We recommend researchers interested in individual differences in temporal structure to first run a reliability study on their paradigm. The importance of reliability has been previously discussed (Hedge et al., 2018; 2020), and has implications for sample size determination.
3. *Formally assess the external validity.* In most of our results, we found that repeatable individual difference in temporal structure failed to show external validity. As we found high positive correlations between temporal structure and performance, and we know there is some within-subject association between MRT performance and subjective ratings of being off-task, it would have been easy to conclude that people who are more off-task show increased temporal structure. Only by including the measure of subjective off-taskness into our design were we able to reject this conclusion. As such, our findings highlight the importance of a multi-modal approach when studying temporal structures, particularly as their neurocognitive mechanisms remain largely mysterious.
4. *Be transparent about analysis choices and their effects.* Small changes in the analysis pipeline may lead to substantial changes, highlighting the importance of transparent reporting. Here we checked our results against two types of analysis choices (frequency length and methods of dealing with missing data). By far, the largest part of the found patterns held up, but there were some obvious exceptions. Transparency also makes it easier to integrate different findings from the literature.
5. *Estimate the autocorrelation.* AC1 may be best suited as a potential biomarker, as its repeatability is high, and the measure is relatively easy to implement in practical settings. Of course, repeatability is not the only construct of interest in neurocognitive variables. Most measures did correlate to performance – and may still be useful for capturing moment-to-moment fluctuations in the data and correlating them to moment-to-moment fluctuations in behavioural and neural data. However, the autocorrelation still comes with some advantages here. Unlike long-term processes, the presence of an autocorrelation can be easily tested with rather unambiguous results, and its estimation is a direct representation of the effect size, with its squared value indicating the amount of explained variance. It also requires fewer analysis choices, and its temporal range can straightforwardly be increased with lag size. Furthermore, the long-range parameters can be difficult to justify if there is no evidence for long-term structure in the RT data. We therefore advocate that the autocorrelation should be included alongside other measures, facilitating comparisons across aims and paradigms.
6. *Manipulate the temporal dependencies.* The underlying mechanisms driving the relationship between performance and temporal structure remain unknown. In particular, it is unclear how the structures behave under different conditions (e.g., different cognitive loads or attentional constraints), and with what kind of neural processes they are associated (as the literature often assumes but rarely empirically tests underlying neural mechanisms), which gets in the way of coming up with clear falsifiable predictions (also see Wagenmakers et al., 2012). Some may argue that the temporal structures should manifest similarly under different conditions – reflecting their ‘ubiquitous nature’ – but this would make the measures mostly uninformative. Similarly, some may argue that healthy participants should exhibit temporal structure largely to the same extent, because all neural systems should have spontaneously converged towards criticality. As noted above, this was clearly not the case in our data. Future research may therefore aim to directly manipulate the temporal structures with different experimental conditions (see Perquin et al., 2020 for one such an attempt) – to get a clearer idea of their neural-cognitive mechanisms.

## Conclusion

The idea that there is meaningful information in what was so far treated as neural or behavioural noise has gained traction over the years, fuelled by growing evidence that spontaneous variability shows temporal structure. In the literature, the emphasis is often put specifically on long-range temporal structures, thought to reflect a universal property of brain-cognition systems. Across new and archival data spanning a variety of sensorimotor and cognitive tasks, we found clear evidence of behavioural temporal structure – and indeed, in many instances the structures were long-term. However, this was not universal across participants and tasks: many RT series only showed short-term dependency. The clarity of temporal structures in behaviour varied systematically across individuals and was, in many tasks, remarkably repeatable. This makes it theoretically possible that they are informative of individual differences in other domains. However, although they were internally related to performance, they were not informative of concomitant attentional state, temporal structure in other tasks or externally assessed attentional traits. Therefore, the glaringly open question remains what they *can* inform us about. Combined with the lack of consistent empirical evidence for a link between behavioural and neural temporal structures, it seems now clear that temporal dependencies are not the unifying manifestation of one overarching stable trait across all circumstances.

## Acknowledgements

We are grateful to Christina Jin and Thomas Anderson for sharing their data with us. We would also like to thank Rachel Draper, Laura Daniells, Laura Fleetwood, Joel Bentley, Catrin McAdams, Megan King, and Eve Evans for their help with collecting the data, and Simon Farrell for sharing his R code for time series analyses.

## Author note

Our own raw data from the reported analyses will be made publicly available upon publication (view-only link for peer review: https://tinyurl.com/hru8rfzy), alongside the analysis code and jasp files.

On behalf of all authors, the corresponding author states that there is no conflict of interest.

## Appendix

Below, we describe the Methods from our own data in detail for both collected cohorts. As both data cohorts have been collected in collaborative projects, including student undergraduate projects, not all measures were analysed for the current research aims. For transparency purposes, we do report all taken measures from the experiments.

Questionnaire scores and eye movement data recorded before and after the task for cohort 1 are published in Perquin & Bompas (2019), alongside other datasets.

This study was not preregistered.

### Metronome Task

#### Participants

Participants were undergraduate Psychology students who participated for course credits. They were specifically excluded from participation if they had a prior clinical diagnosis of ADHD or depression, if they did not have normal or corrected-normal hearing, or if they had significant difficulties with fine motor control. Participants were instructed to refrain from alcohol and drugs in the 24 hours prior to the experiment. The study was approved by the local ethics commission. Combined over the two cohorts, we had analysable datasets from 139 participants, with 73 having done both MRT sessions (see section *Data preparation and analysis* for details).

##### Cohort 1

84 healthy participants (69 female, 14 male, one other, aged 18-25) participated in a laboratory experiment. One was excluded because of technical issues. Of these, 24 participants performed the behavioural task twice.

##### Cohort 2

81 healthy participants (62 female, 18 male, 1 preferred not to say, aged 18-21) participated in an online video conferencing experiment. Three were excluded during screening for having a diagnosis of depression, five had incomplete data, three dropped out before the behavioural session, and for another three no behavioural responses were recorded during the MRT. This relatively high attrition rate is likely due to the online nature. In total, this resulted in data from 66 participants, of whom 61 performed the behavioural session twice.

#### Materials

##### Cohort 1

The behavioural paradigm was generated on a Viglen Genie PC with MATLAB version 8 (The Mathworks, Inc, Release 2015b) and Psychtoolbox-3 (Brainard, 1997; Kleiner et al., 2007; Pelli, 1997), and was displayed on an ASUS VG248 monitor with a resolution of 1920 by 1080 and a refresh rate of 144 Hz. The background was light grey throughout the experiment, with the fixation point and text in white. During the MRT task(s), eye movements and pupil dilation were recorded with an Eyelink 1000 (SR Research), with participants seated with their head in a chinrest to limit motion (at 615 cm distance from the screen).

##### Cohort 2

Due to the COVID-19 pandemic, data were collected online. The experiment was generated in Psychopy 3 and was run using Pavlovia (Pierce et al., 2019). Participants performed the experiment on their personal devices. The background was light grey throughout the experiment, with the fixation point and text in white.

#### Questionnaires

##### Cohort 1

Participants completed the Adult ADHD Self-Report Scale (ASRS-v1.1; Kessler et al., 2005). This scale consists of eighteen items on a scale from 0 (*“Never”)* to 4 (*“Very often”),* and is composed of two subscales: Inattention and Hyperactivity / impulsivity (Kessler et al., 2005; Reuter, Kirsch & Hennig, 2006). Internal consistency of the ASRS-v1.1 is high (Cronbach’s ranged from .88-.94; Adler et al., 2006; 2012).

To measure mind wandering tendencies in daily life, participants completed the Daydreaming Frequency Scale (DFS; Singer & Antrobus, 1963), a subscale of the Imaginal Processes Inventory that consists of twelve 5-point items. The DFS also has a high internal consistency, as well as high test-rest reliability (Cronbach’s α = .91, test-retest reliability with interval of maximum one year = .76; Giambra, 1980).

Furthermore, participants filled in the UPPS-P Impulsive Behaviour Scale (Lynam, Smith, Whiteside & Cyders, 2006; Whiteside & Lynam, 2001). This questionnaire consists of 59 items, scored on a scale from 1 (*“agree strongly”*) to 4 (*“disagree strongly”*), and is composed of five subscales: positive urgency, negative urgency, (lack of) premeditation, (lack of) perseverance, and sensation seeking.

All participants also filled in six other questionnaires, which were not analysed in the current study: the Beck Anxiety Inventory Second edition (Beck & Steer, 1993), Beck Depression Inventory Second edition (Beck, Steer & Brown, 1996), Short form Wisconsin Schizotypy scales (Winterstein et al., 2011), Five-facet Mindfulness Questionnaire (Baer, Smith, Hopkins, Krietemeyer & Toney, 2008), Toronto mindfulness scale (Lau et al., 2006), and Positive and Negative Affect Schedule (Watson, Clark & Tellegen, 1988).

##### Cohort 2

Participants completed the ASRS-v1.1 (Kessler et al., 2005). For the purpose of other studies, participants also completed the Beck Depression Inventory Second edition (Beck et al., 1996) and questions related to covid-related social isolation and sleep quality.

#### Procedures

##### Cohort 1

Participants came to the lab for one session. After eye tracker calibration, participants took part in a four-minute resting state session (eyes open), to get them into a common baseline state before starting the behavioural task. Next, they performed the MRT (~25 minutes). After the task, they performed another resting state session (eyes open), and then filled in questionnaires. In total, this took about 1.5 hours. Of the 83 participants, 25 of them then performed the MRT again, after watching one of two video clips of 3 and 5 min.

Figure 2 shows an overview of the MRT task over time (Seli et al., 2013). Each trial lasts 1300 ms. In the middle of the trial (650 ms after onset), a short tone is presented to the participants (~75ms). The task of the participant is to press *on* this tone, such that perfect performance is indicated by a complete synchrony between tones and presses. Rhythmic reaction time (RT) is measured as the relative time from the press to the tone (with RT = 0 being a perfect response). The RT series throughout the task were used to measure performance and temporal structures in performance.

Throughout the task, participants were presented with thought probes. The first question related to their subjective rating of attention just prior to the thought probe appeared (“Please record a response from 1 to 9 which characterises how on task you were just before this screen appeared”, with 1 as ‘completely ON task’ and 9 as ‘completely OFF task’). Based on their rating, they then received five follow-up questions related to the content, temporality, valence, and intentionality of their on-task (if their rating was 1-3) or off-task (if their rating was > 3) thoughts, as well as to their motivation. The follow-up questions were not analysed in the current research. In total, the experimental phase of the MRT consisted of 21 blocks with 50 trials each (1050 trials in total), with one thought probe in every block. Probes were presented pseudo-randomly. To make sure the probes did not follow too closely after each other, they were never administered in the first five trials of a block.

The random lottery reward system (Cubitt, Starmer & Sugden, 1998) was used to motivate participants to keep up good performance throughout the task. After the session, one trial *n* was randomly extracted, and if the standard deviation of trial *n* to trial *n-4* was below .075 (indicating consistent performance in that time window), the participant received a reward of £5. The cut-off of .075 was based on pilot data, chosen such that ~20% of the participants would receive the reward.

Before the experimental phase of the MRT, participants received a training block of 50 trials to learn the rhythm of the tone. At training trial 15, they were presented with a thought probe. After the training, the participant received feedback on their performance from the experimenter, to make sure they understood the task. Participants were also told how many of their trials would qualify for the reward, to provide them motivation to keep up good performance.

##### Cohort 2

Participants attended one online session through video conferencing. The session started with a brief plenary explanation from the experimenter. They first received a Qualtrics link, in which they completed informed consent and filled in the ASRS.

Next, they performed the MRT (~25 minutes). The trials, blocks, and probes were kept equal to the MRT of the first cohort. The only differences were the follow-up questions, which were also not analysed in the current research, and the absence of a lottery-based rewards system. After the first behavioural task, participants filled in a short questionnaire related to sleepiness, and then were instructed to have a break in-between the two tasks. On average, they completed the second task 30 minutes after the first task.

#### Data preparation and analysis

For each participant, the total percentage of omissions was calculated, and participants with more than 10% omissions were excluded from analyses (following the procedure of Seli et al., 2013). Four participants from cohort 1 and five participants from cohort 2 were excluded due to having too many omissions in the first session (~6% of total useable datasets) and furthermore, one participant was excluded for responding in anti-phase with the tone. Data from another 6 participants (one from cohort 1) were excluded from the second MRT session due to too many omissions (~7% of total useable datasets).

Performance was quantified on the basis of each RT series using the standard deviation of the RT (SD, reflecting consistency), as is common in studies using the MRT. The first five trials as well as the five trials following each thought probe were excluded from SD calculation. Two measures of subjective attentional state were calculated on each thought probe rating series: the mean and SD of the 21 ratings. Independent two-sample t-tests were conducted on SD and mean subjective attentional rating to see if systematic differences appeared between the two cohorts. As the evidence favoured an absence of difference (BF_01_ = 2.2, 3.5, 3.8, and 4.5 respectively for the two measures on the first and the second MRT sessions), data of the two cohorts were combined across all analyses to increase statistical power. Still, as minor differences between cohorts might contaminate the intra- and inter-individual correlation analyses, RT series were normalised for each participant across the two cohorts, using Y(c,n) = X(c,n) – mean(X(c,1):X(c,Nc)), where *Y* is the normalised data series, *X* is the original RT series, *c* is cohort, *n* is participant number and *Nc* is the number of participants in each cohort). As the distribution of SD was not normal on the group level, it was log-transformed (decreasing its Skewness value from 2.0 to .6, and its Kurtosis from 5.4 to .5).

The mean and SD of subjective ratings were also correlated (r = .26, BF_10_ = 12.6), verifying the assumption that participants who report being more off-task also switch more between ratings on average, though this relationship is not very strong. Across individuals, there was no evidence that the subjective measures correlated with any of the objective measures (r ranging from 0.05 to 0.15, BF_01_ ranging 2.2 to 8.2). For completion, we also report the standard within-subject analysis (as per Anderson et al., 2021; Laflamme et al., 2018; Seli et al., 2013): variance was calculated on the last five trials before each probe, and its logarithmically transformed value was correlated within participants to the attentional state ratings. Indeed, increased levels of being off task were associated with increased variability in the RTs (median Kendall’s τ = .08, BF_10_ > 1000). We report this here—rather than in the main results — as this was not the purpose of the present study. We note that, consistent with previous studies, local correlations between performance and attention ratings are highly significant, but their shared variance is weak, consistent with the idea that most of the variability in performance escapes consciousness (see Perquin et al., 2020 for empirical implications of this idea).

### Archival datasets

Here we describe the Methods from the archival datasets. As these Methods have been described in their original papers, we will only summarise the key points and describe our current data analyses. These archival datasets were considered after having fully analysed the results from our own data. None allowed to test all our questions, but each of them provided an opportunity to test the generalisability of some of our conclusions. All datasets analysed are included in this report.

#### MRT: Anderson et al. (2021)

We received data from 375 participants (all with omissions rates lower than 10%), who had participated in a single-session laboratory study on the MRT. The timing of the MRT was the same as our design (1300 ms), with one thought probe being presented once in each 50-trial block. Participants completed 18 blocks (compared to our 21), giving 900 trials in total. On the thought probes, they were asked to categorise their subjective attentional state using the following six response options: 1) completely on-task, 2) mostly on-task, 3) mostly mind-wandering unintentionally, 4) completely mind-wandering unintentionally, 5) mostly mind-wandering intentionally, and 6) completely mind-wandering intentionally. After the MRT, participants completed the Attention-Related Cognitive Errors Scale (ARCES; Cheyne et al., 2006).

The ARCES consists of 12 questions, aiming to measure individuals’ tendency to experience attentional lapses in daily life (e.g., *“I have absent-mindedly placed things in unintended locations (e.g., putting milk in the pantry or sugar in the fridge).”*). It has a high internal consistency, with Cronbach’s α = .88 (Cheyne et al., 2006).

##### Data preparation and analysis

Six participants were excluded based on qualitative checks by the original paper (Anderson et al., 2021). Furthermore, 17 participants were excluded by us for responding in antiphase with the tone and/or having pronounced bimodal RT distributions – leaving us with 352 participants for analysis. Again, we calculated the SD of RT for each participant.

However, group distributions of RT revealed two noteworthy differences between these and our own MRT data. Firstly, mean RT (calculated over the raw RT series, with a possible range of −.65 to .65, and reflecting bias compared to the tone) was high on average (.22 s), indicating that participants were not responding *on* the tone as per instructions, but rather as a response to it. In comparison, the group means in our data from the first session were −.06 and −.09 s respectively for the first and second cohort, indicating that participants were responding anticipatory to the tone. Secondly, SD did not correlate to the mean of the absolute RT (|RT|, calculated over the absolute values of the RT series, with a possible range of 0 to .65, and reflecting absolute distance to the tone; r = .01, BF_01_ = 14.4). In contrast, in our MRT data, SD and mean |RT| were highly correlated (r = .75, log(BF_10_) = 53.8) – indicating that participants who performed well pressed close to the tone and did so consistently, while poorer-performing participants had larger asynchronies and were more variable. This is more in line with what one would expect from task instructions, which would lead well-performing participants to *anticipate* the tone rather than respond to it. We note that, the performance measures from this dataset are thus more difficult to interpret on a between-subject level than our MRT data.

Mean and SD of the subjective ratings (scaled 1-6) were calculated for each subject. These were positively correlated to each other (r = .49, log(BF_10_) = 44.7). As a sanity check, we also ran all analyses using the percentage of being off-task (number of off-task (3 to 6) responses / total number of responses) instead of the mean rating. These did not show any different result patterns. None of the performance measures correlated with the subjective ratings across participants (*r* ranging .03-.11, BF_01_ ranging 2.1-12.7). On a within-subject level, the attentional state ratings correlated with increased within-subject variability of RT (quantified as SD on the last five RT before the thought probes, as reported in the original paper

AC1, PSD slope, and ARFIMA parameters were calculated as above. Settings for estimating the Bayes Factors were also kept the same. DFA was performed over non-overlapping blocks log-linearly spaced from a minimum of 4 trials to 256 trials.

#### Cognitive Control: Hedge et al. (2018)

Here we describe three experiments that were conducted for a reliability study of task performance in a battery of well-known cognitive paradigms (Hedge et al., 2018). As in the original paper, we have combined the data from Experiment 1 and 2, as these included the same tasks.

##### Experiment 1 and 2

107 participants took part in two sessions (Experiment 1: 50, Experiment 2: 62) in a laboratory experiment. Nine participants were excluded from analysis on all four tasks, and another five and one only on the Stop-signal and Stroop task respectively, based on poor performance as per the original paper.

Participants performed four tasks of ~20 minutes each. The first was an Eriksen Flanker task, in which participants have to discriminate the direction of an arrow. The task consisted of three conditions (congruent: flanker arrows pointed in the same direction as target arrow, incongruent: flanker arrows pointed in the opposite direction as target arrow, and neutral: flankers were lines rather than arrows), of which participants completed 240 trials for each (720 trials in total). The second was a Stroop task, in which participants had to name the colour of the word. The task consisted of three conditions (congruent: the written word was the same as the physical colour of the word, e.g., the word ‘red’ written in red letters, incongruent: the written word was a different colour as the colour, and neutral: the written word was a non-colour word), of which participants completed 240 trials for each (720 trials in total, ~20 minutes). The third task was a Stop-Signal task, in which participants had to respond if the presented stimulus was a square or a circle. On 25% of the tasks, a tone (stop signal) was presented indicating to withhold their response (with tone latency from the stimulus onset being an individual adaptive measure, as is common with this task). Participants performed 600 trials in total (25% stop-trials). The fourth task was a Go/No-Go task. In each block, participants were presented with one of four letter stimuli. They were instructed to respond as quickly as possible to three of the letters (75% go trials) and withhold a response to the fourth (25% no-go trials). Participants performed 600 trials in total.

The order of the tasks was counterbalanced across participants. Participants completed each of the four tasks in a first session. Three weeks later, they returned to the lab and completed all four tasks again.

##### Experiment 3

40 participants took part in two sessions in a laboratory experiment. They performed three tasks, each of ~20 minutes. The first was the Posner Cuing Task, in which participants were presented with a stimulus on the left or the right side of the screen, and had to press a button as soon as they detected it. Before stimulus onset, they were presented with a cue (SOA = 300-600 ms, in steps of 100 ms), which was accurate in 80% of trials. The second task was a NAVON task, in which participants were presented with a large ‘S’ or a large ‘H’, which consisted of smaller ‘S’ and ‘H’ letters. These refer respectively to the global and local level of the stimulus, which could either be congruent or incongruent with each other. On half the trials, participants had to identify the global level, and on the other half, the local level. The third task was the Spatial-Numerical Association of Response Codes (SNARC) task, in which participants had to respond if a presented letter was smaller or larger than 5. On half of the trials, the ‘smaller’ response had to be indicated by a left-sided button press and the ‘bigger’ response by a right-sided button press. On the other half of trials, these response mappings were reversed.

Participants completed 640 trials in total for each task. Task order was counterbalanced across participants. Participants completed all three tasks in a first session. Three weeks later, they returned to the lab and completed all tasks again.

##### Data preparation and analysis

For each of the seven tasks and for each session separately, the AC1, PSD slope, and ARFIMA parameters were calculated as above on the RT series. Settings for estimating the Bayes Factors were kept the same. DFA was performed over non-overlapping blocks log-linearly spaced from a minimum of 4 trials to 256 trials.

#### Visual Search and SART (Jin et al., 2019)

30 participants took part in an EEG study, in which they completed two tasks. The first was a Sustained Attention to Response Task (SART), which has been used extensively to study subjective attentional states (e.g., Christoff et al., 2009; McVay & Kane, 2009; Qin et al., 2011; Robison et al., 2019; Smallwood et al., 2008; 2009; Stawarczyk, Majerus & D’Argembeau, 2013; Van Vugt & Broers, 2016). Participants had to respond with a button press if the stimulus was a lower-case word (frequent case, 89%) and had to withhold their response if the stimulus was an upper-case word (infrequent case, 11%). They completed 378-486 trials. The second task was a Visual Search task, in which participants had to indicate if a target was present or absent (with targets being present on 50% of trials). They completed 420 trials.

In both tasks, participants were occasionally presented with a thought probe (presented every 7-24 trials, with a total of 54 probes per task for each participant). They were asked to categorise their attentional state using one of the following six response options: 1) “I entirely concentrated on the ongoing task”, 2) “I evaluated aspects of the task (e.g., my performance or how long it takes)”, 3) I thought about personal matters”, 4) I was distracted by my surroundings (e.g., noise, temperature, my physical condition”, 5) I was daydreaming, thinking of task unrelated things, or 6) “I was not paying attention, but my thought wasn’t anywhere specifically”.

##### Data preparation and analysis

As the RT series contained some extreme outliers (particularly in the Visual Search task), RTs that were 3 SD above the mean were first excluded for each participant. In the SART, .07 % of the trials were excluded on average (range: 0-2.1%), and in the Visual Search, 1.1 % of the trials were excluded (range: 0-2.9%).

The original paper reported relationships between the subjective ratings and performance. As the categories from the attentional state ratings do not correspond directly to depth, we instead calculated the percentage off-task responses for each participant (number of 3 to 6 responses / total number of responses * 100%).

AC1, PSD slope, and ARFIMA parameters were calculated on the RT series for both tasks separately. Settings for estimating the Bayes Factors were kept the same. DFA was performed over non-overlapping blocks log-linearly spaced from a minimum of 4 trials to 128 trials.

#### RT tasks (Wagenmakers et al., 2004)

We also reanalysed the data from Wagenmakers et al. (2004), an important milestone in the temporal dependencies literature, containing six participants, who performed two simple RT, two choice RT, and two time-estimation tasks. This data did not lend itself to our specific research questions, as the sample size is too small to estimate reliable correlation coefficients, but has been plotted in Figure 3 alongside the rest of the data to compare the weight of the measures.

Six participants took part in a behavioural study investigating the presence of 1/f noise in RT series, in which they performed simple RT, choice RT, and time estimation tasks. One each trial, they were shown a number from 1-9. In the simple RT task, they had to press a button as soon as they saw the stimulus. In the choice RT task, they had to discriminate whether the number was odd or even. In the time estimation task, they had to respond when they thought one second had passed since stimulus onset.

For each task, participants performed a ‘short’ and a ‘long’ version, relating to the time between response and next trial (response-to-stimulus interval; RSI). The original paper found no significant differences in temporal structure within tasks between short and long RSI. Each participant performed 1024 trials for all six tasks.

##### Data preparation and analysis

AC1, PSD slope, and ARFIMA parameters were calculated on the RT series for each tasks separately. Settings for estimating the Bayes Factors were kept the same. DFA was performed over non-overlapping blocks log-linearly spaced from a minimum of 4 trials to 256 trials.

## Supplementary Materials A

**Table A1.**
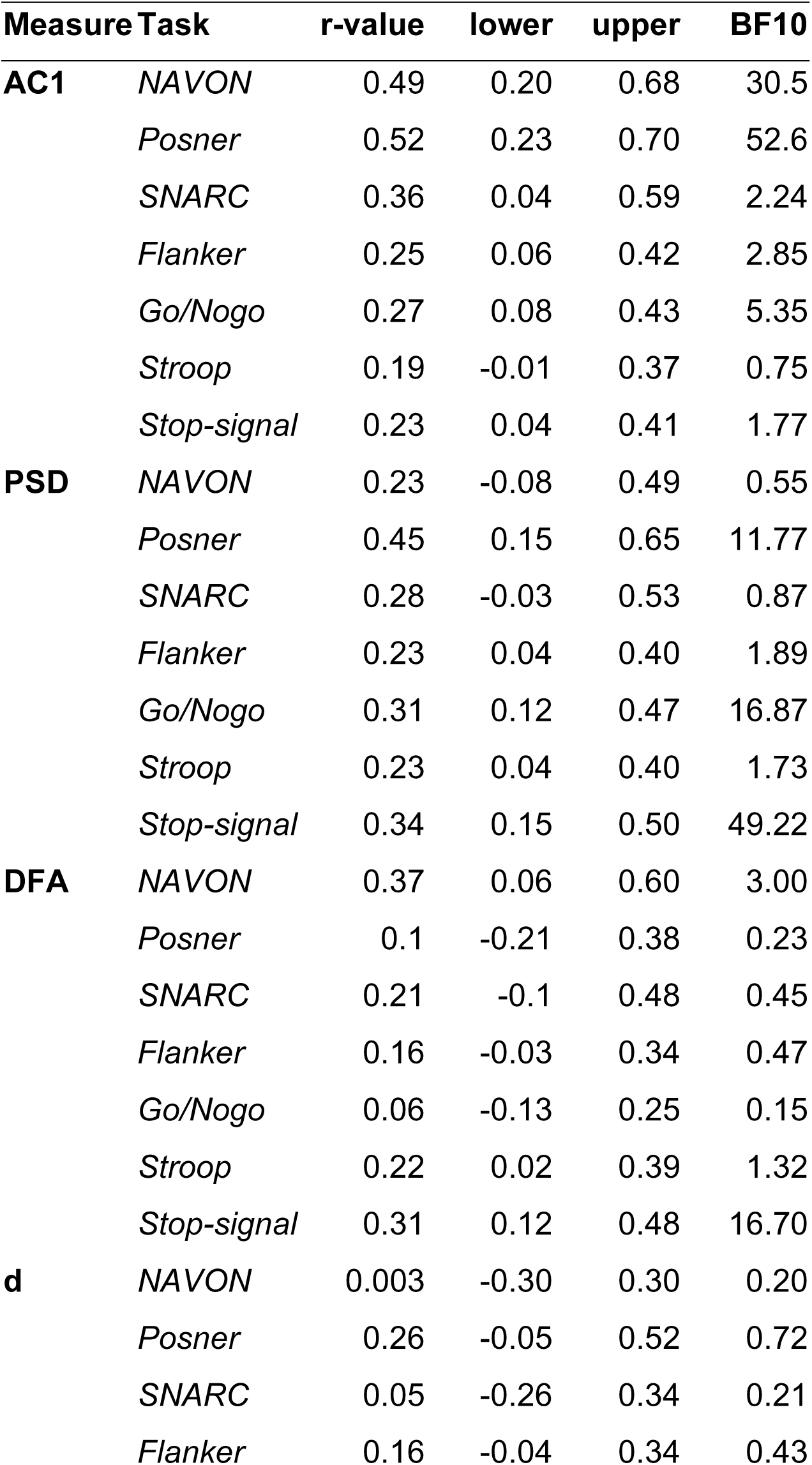
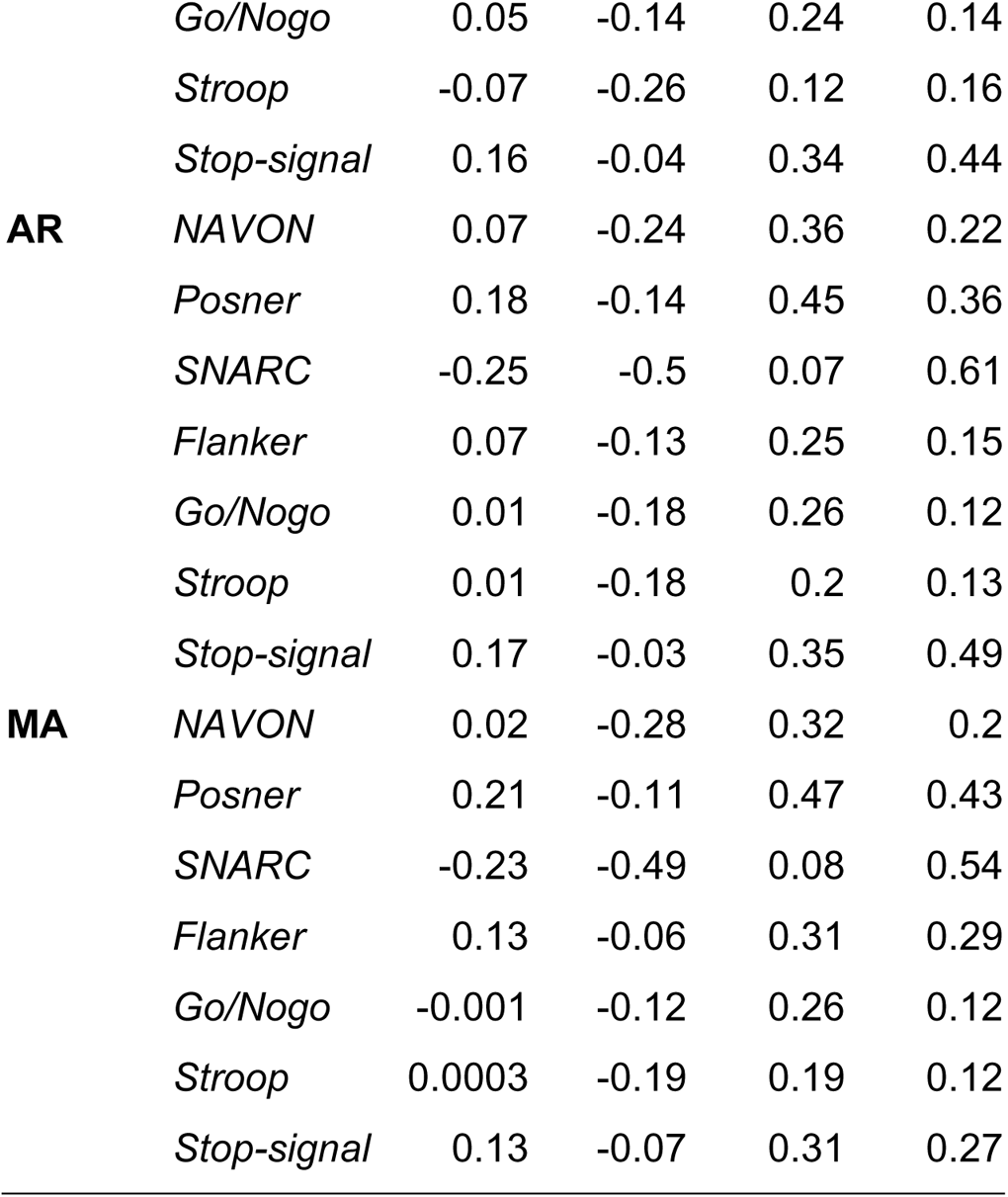
Full list of the correlation coefficients (r-value) with corresponding 95% CI ([lower, upper]) and Bayes Factor (BF10) for the within-task repeatability of cognitive tasks, as displayed in Figure 6.

**Table A2.**
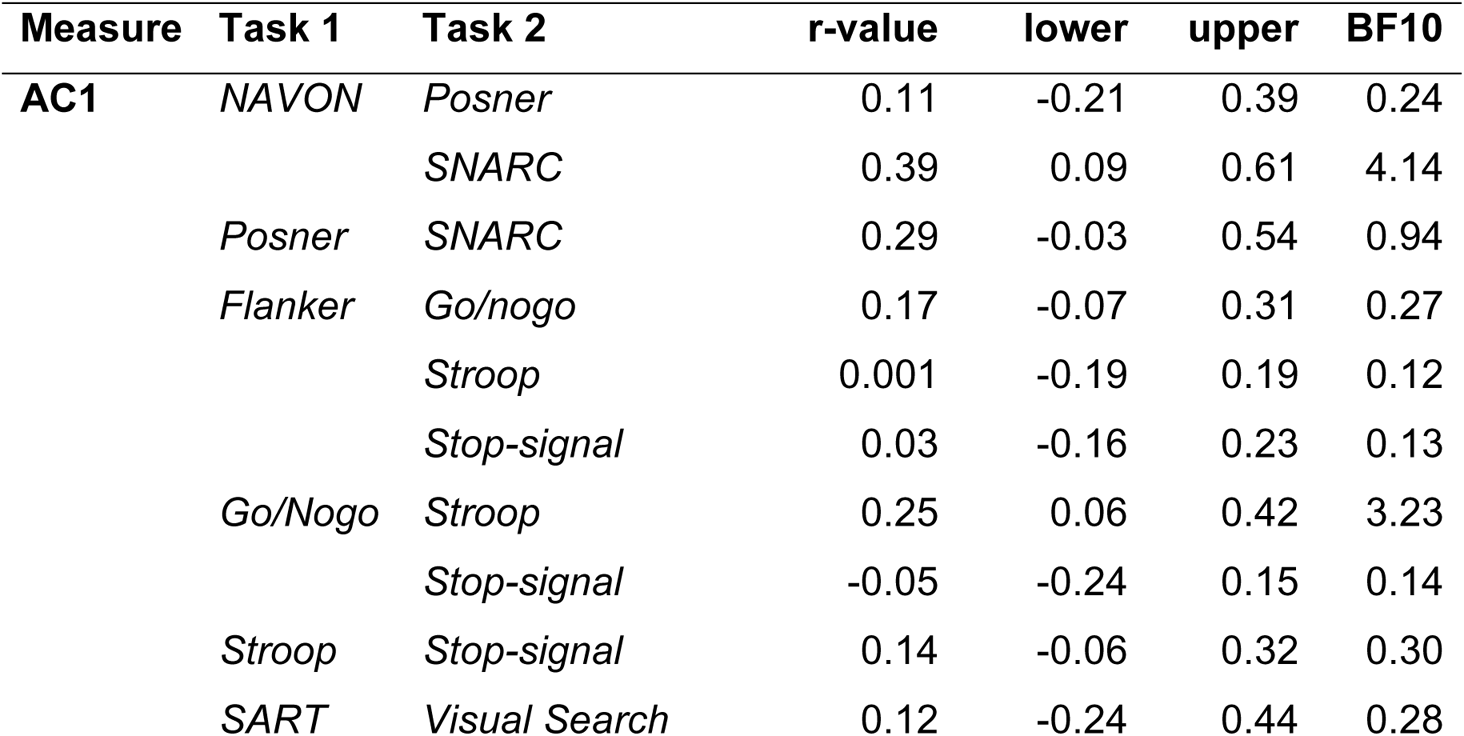
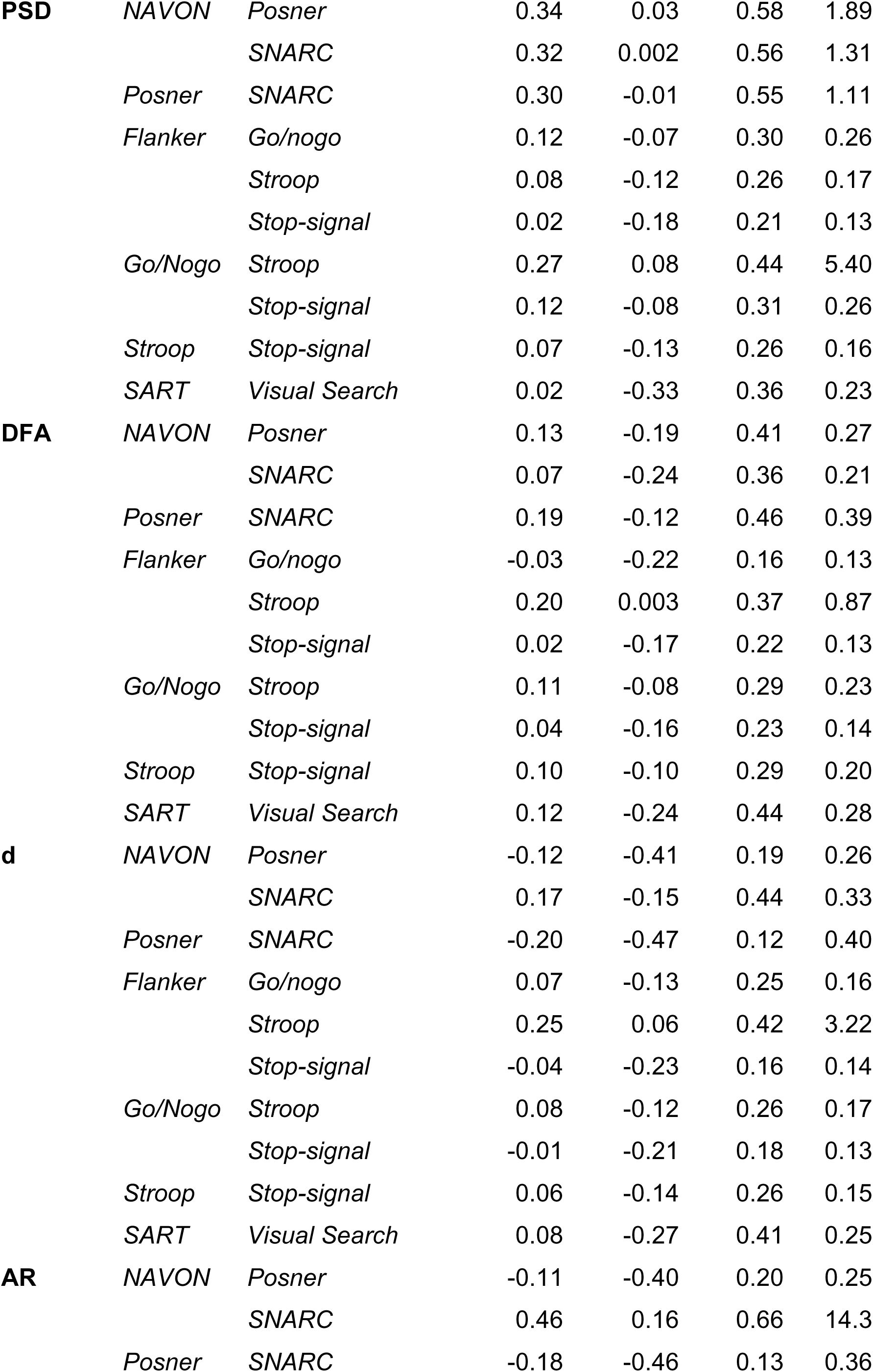
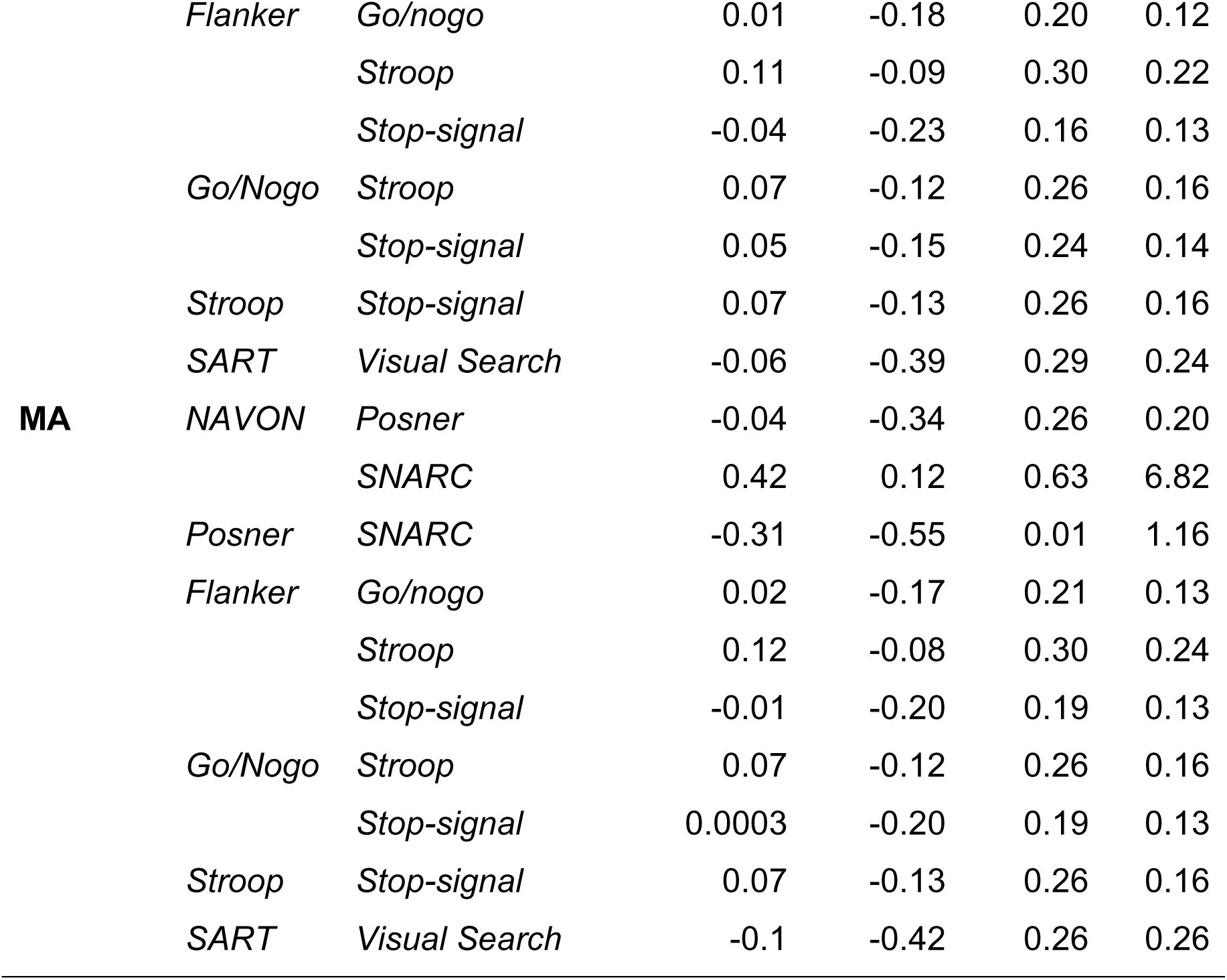
Full list of the correlation coefficients (r-value) with corresponding 95% CI ([lower, upper]) and Bayes Factor (BF10) for the between-task repeatability of cognitive tasks, as displayed in Figure 6.

## Supplementary Materials B

**Table B.** Pearson r-values between the different measures of temporal dependency on each of the archival datasets, with each row reflecting one correlation pair. Green and red fonts indicate clear evidence against and for the null respectively; black font indicates no reliable conclusion can be drawn.

|  | MRT | SART | VS | Flanker | Stroop | Stopsignal | Go/No-go | NAVON | SNARC | Posner |
| --- | --- | --- | --- | --- | --- | --- | --- | --- | --- | --- |
| AC1-PSD | .93 | .92 | .87 | .89 | .88 | .76 | .89 | .77 | .91 | .85 |
| AC1-DFA | .77 | .66 | .55 | .60 | .69 | .68 | .67 | .86 | .57 | .59 |
| AC1-d | .04 | .23 | .16 | .31 | .37 | .50 | .33 | -.26 | .16 | .41 |
| AC1-AR | .26 | -.09 | .07 | -.02 | -.10 | .13 | .18 | .20 | -.01 | -.18 |
| AC1-MA | .07 | -.17 | -.07 | -.12 | -.12 | .13 | .12 | .04 | -.07 | -.19 |
| PSD-DFA | .69 | .62 | .59 | .57 | .68 | .72 | .64 | .64 | .56 | .65 |
| PSD-d | -.02 | .22 | .09 | .22 | .23 | .31 | .28 | -.41 | .03 | .32 |
| PSD-AR | .22 | -.17 | .10 | .03 | .19 | .02 | .14 | .18 | -.07 | -.18 |
| PSD-MA | -.01 | -.26 | -.02 | -.06 | .22 | .04 | .07 | -.03 | -.15 | -.20 |
| DFA-d | .10 | .32 | .16 | .48 | .33 | .59 | .46 | -.19 | .51 | .57 |
| DFA-AR | .19 | -.04 | -.07 | -.16 | -.15 | -.04 | .18 | -.02 | -.25 | -.12 |
| DFA-MA | .12 | -.04 | -.11 | -.15 | -.13 | .04 | .20 | -.14 | -.18 | -.04 |
| d-AR | -.59 | .03 | -.44 | -.55 | -.18 | .21 | -.25 | -.41 | -.07 | -.02 |
| d-MA | -.27 | .22 | -.27 | -.39 | .03 | .41 | -.08 | -.12 | .13 | .22 |
| ar-MA | .94 | .97 | .97 | .96 | .97 | .97 | .98 | .95 | .97 | .96 |

## Supplementary Materials C

**Table C1.**
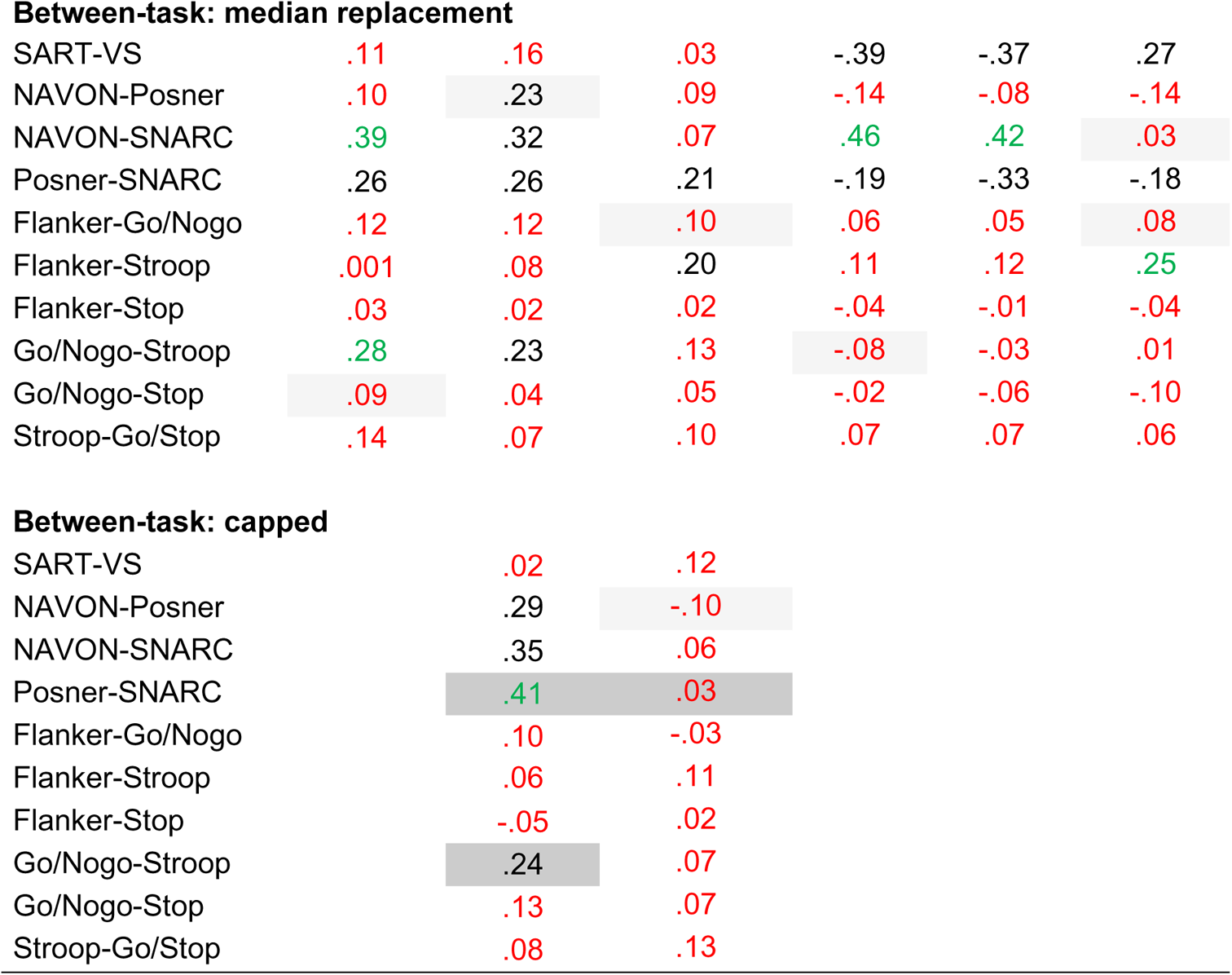
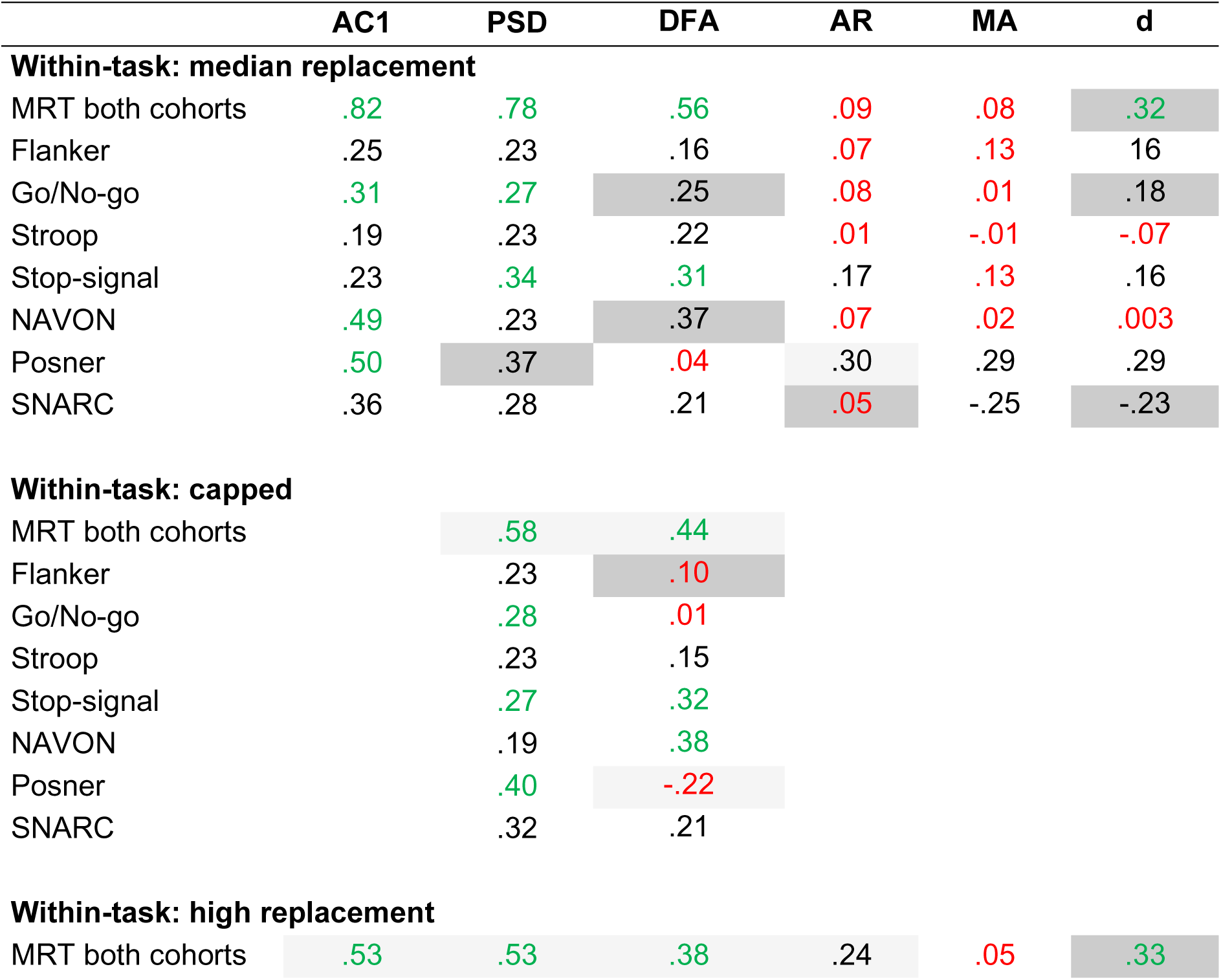
Within- and between-task repeatability analyses using different analysis choices, in which the missing values in the RT series were replaced either by the median value or highest possible (MRT only) value, or in which the frequency was capped (PSD and DFA only). Shown are the r-values of the correlation analyses, with green and red fonts indicating clear evidence against and for the null respectively. R-values which Bayesian evidence has changed evidence are highlighted in dark grey, and r-values which evidence did not change but were at least .10 higher or lower than the original estimate are highlighted in light grey.

|  | AC1 | PSD | DFA | AR | MA | d |
| --- | --- | --- | --- | --- | --- | --- |
| **Within-task: median replacement** |  |  |  |  |  |  |
| MRT both cohorts | .82 | .78 | .56 | .09 | .08 | .32 |
| Flanker | .25 | .23 | .16 | .07 | .13 | .16 |
| Go/No-go | .31 | .27 | .25 | .08 | .01 | .18 |
| Stroop | .19 | .23 | .22 | .01 | -.01 | -.07 |
| Stop-signal | .23 | .34 | .31 | .17 | .13 | .16 |
| NAVON | .49 | .23 | .37 | .07 | .02 | .003 |
| Posner | .50 | .37 | .04 | .30 | .29 | .29 |
| SNARC | .36 | .28 | .21 | .05 | -.25 | -.23 |
| **Within-task: capped** |  |  |  |  |  |  |
| MRT both cohorts |  | .58 | .44 |  |  |  |
| Flanker |  | .23 | .10 |  |  |  |
| Go/No-go |  | .28 | .01 |  |  |  |
| Stroop |  | .23 | .15 |  |  |  |
| Stop-signal |  | .27 | .32 |  |  |  |
| NAVON |  | .19 | .38 |  |  |  |
| Posner |  | .40 | -.22 |  |  |  |
| SNARC |  | .32 | .21 |  |  |  |
| **Within-task: high replacement** |  |  |  |  |  |  |
| MRT both cohorts | .53 | .53 | .38 | .24 | .05 | .33 |
| **Between-task: median replacement** |  |  |  |  |  |  |
| SART-VS | .11 | .16 | .03 | -.39 | -.37 | .27 |
| NAVON-Posner | .10 | .23 | .09 | -.14 | -.08 | -.14 |
| NAVON-SNARC | .39 | .32 | .07 | .46 | .42 | .03 |
| Posner-SNARC | .26 | .26 | .21 | -.19 | -.33 | -.18 |
| Flanker-Go/Nogo | .12 | .12 | .10 | .06 | .05 | .08 |
| Flanker-Stroop | .001 | .08 | .20 | .11 | .12 | .25 |
| Flanker-Stop | .03 | .02 | .02 | -.04 | -.01 | -.04 |
| Go/Nogo-Stroop | .28 | .23 | .13 | -.08 | -.03 | .01 |
| Go/Nogo-Stop | .09 | .04 | .05 | -.02 | -.06 | -.10 |
| Stroop-Go/Stop | .14 | .07 | .10 | .07 | .07 | .06 |

**Between-task: capped**
|  |  |  |
| --- | --- | --- |
| SART-VS | .02 | .12 |
| NAVON-Posner | .29 | -.10 |
| NAVON-SNARC | .35 | .06 |
| Posner-SNARC | .41 | .03 |
| Flanker-Go/Nogo | .10 | -.03 |
| Flanker-Stroop | .06 | .11 |
| Flanker-Stop | -.05 | .02 |
| Go/Nogo-Stroop | .24 | .07 |
| Go/Nogo-Stop | .13 | .07 |
| Stroop-Go/Stop | .08 | .13 |

**Table C2.**
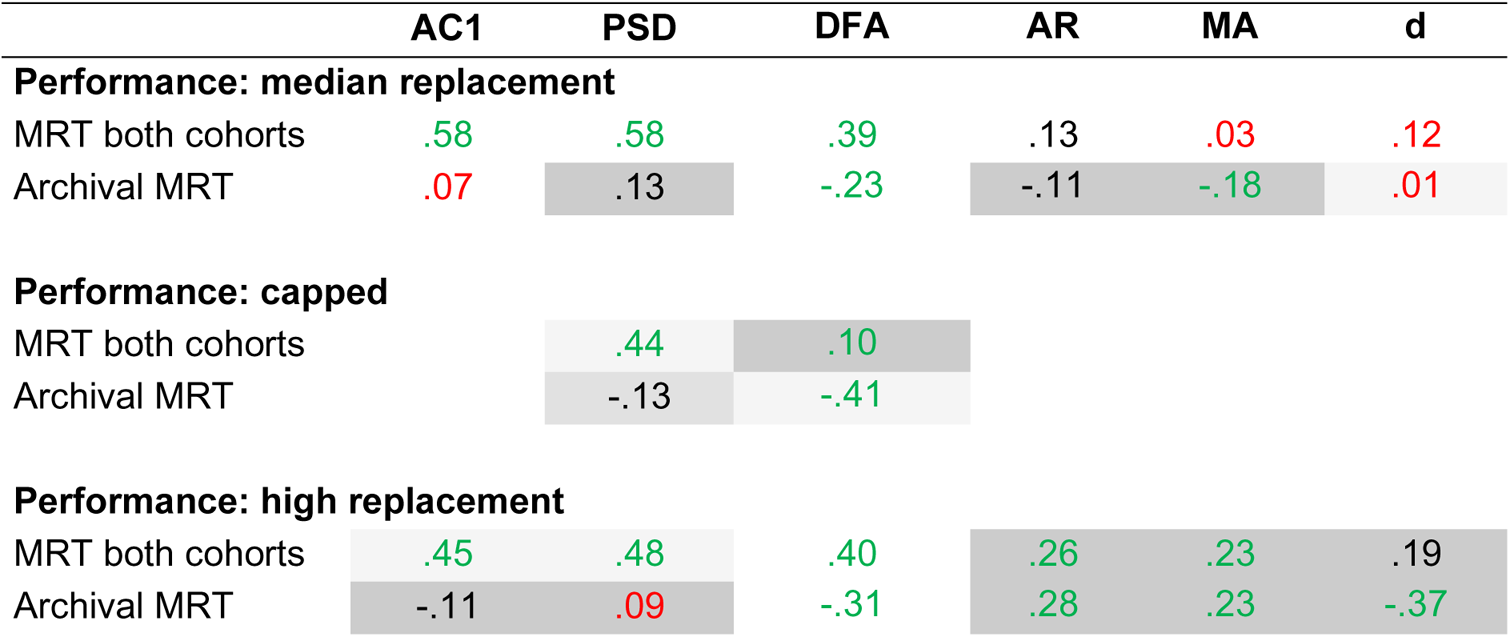
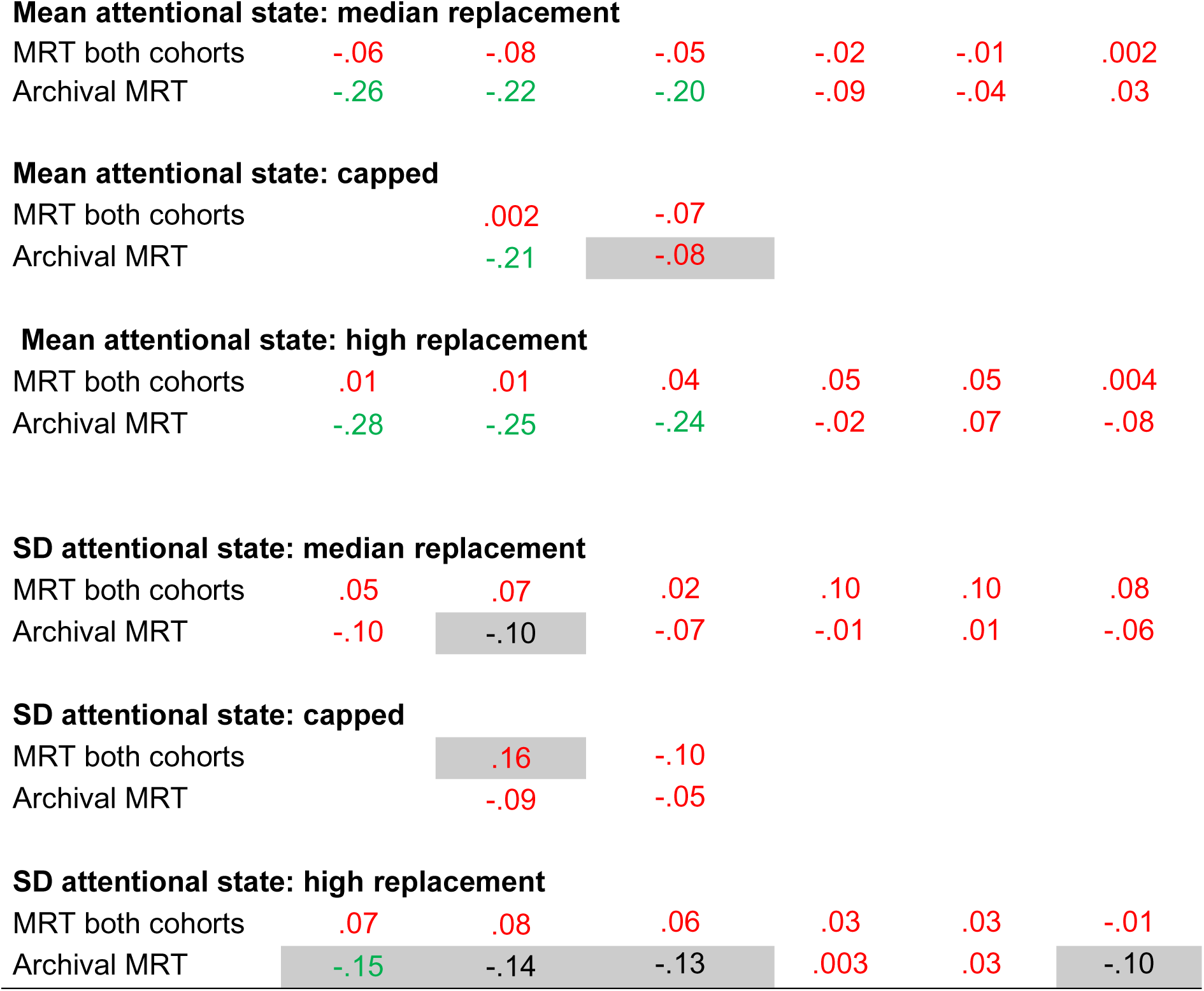
Between-subject correlation analyses between temporal dependency and task measures from the MRT using different analysis choices. Conventions are the same as in Table B2.

|  | AC1 | PSD | DFA | AR | MA | d |
| --- | --- | --- | --- | --- | --- | --- |
| **Performance: median replacement** |  |  |  |  |  |  |
| MRT both cohorts | .58 | .58 | .39 | .13 | .03 | .12 |
| Archival MRT | .07 | .13 | -.23 | -.11 | -.18 | .01 |
| **Performance: capped** |  |  |  |  |  |  |
| MRT both cohorts |  | .44 | .10 |  |  |  |
| Archival MRT |  | -.13 | -.41 |  |  |  |
| **Performance: high replacement** |  |  |  |  |  |  |
| MRT both cohorts | .45 | .48 | .40 | .26 | .23 | .19 |
| Archival MRT | -.11 | .09 | -.31 | .28 | .23 | -.37 |
| **Mean attentional state: median replacement** |  |  |  |  |  |  |
| MRT both cohorts | -.06 | -.08 | -.05 | -.02 | -.01 | .002 |
| Archival MRT | -.26 | -.22 | -.20 | -.09 | -.04 | .03 |

**Mean attentional state: capped**
|  |  |  |
| --- | --- | --- |
| MRT both cohorts | .002 | -.07 |
| Archival MRT | -.21 | -.08 |
| **Mean attentional state: high replacement** |  |  |  |  |  |  |
| MRT both cohorts | .01 | .01 | .04 | .05 | .05 | .004 |
| Archival MRT | -.28 | -.25 | -.24 | -.02 | .07 | -.08 |
| **SD attentional state: median replacement** |  |  |  |  |  |  |
| MRT both cohorts | .05 | .07 | .02 | .10 | .10 | .08 |
| Archival MRT | -.10 | -.10 | -.07 | -.01 | .01 | -.06 |

**SD attentional state: capped**
|  |  |  |
| --- | --- | --- |
| MRT both cohorts | .16 | -.10 |
| Archival MRT | -.09 | -.05 |
| **SD attentional state: high replacement** |  |  |  |  |  |  |
| MRT both cohorts | .07 | .08 | .06 | .03 | .03 | -.01 |
| Archival MRT | -.15 | -.14 | -.13 | .003 | .03 | -.10 |

**Table C3.** Between-subject correlation analyses between temporal dependency and questionnaire scores using different analysis choices. Conventions are the same as above.

|  | AC1 | PSD | DFA |
| --- | --- | --- | --- |
| **Median replacement** |  |  |  |
| ADHD | .04 | .01 | .06 |
| DFS | -.03 | -.08 | .06 |
| UPPS-P | -.02 | -.06 | -.02 |
| ARCES | -.03 | -.01 | -.03 |
| **Capped** |  |  |  |
| ADHD |  | -.04 | -.06 |
| DFS |  | -.04 | -.06 |
| UPPS-P |  | -.09 | .06 |
| ARCES |  | .002 | -.09 |
| **High replacement** |  |  |  |
| ADHD | -.04 | -.03 | .02 |
| DFS | .01 | -.05 | -.02 |
| UPPS-P | -.07 | -.11 | -.09 |
| ARCES | -.07 | -.07 | -.09 |

